# Tolerance to the antifungal drug fluconazole is mediated by tuning cytoplasmic fluidity

**DOI:** 10.64898/2025.12.03.691994

**Authors:** E. Plumb, A. Serrano, L. Chevalier, J. Elferich, L.R. Sinn, N. Grigorieff, M. Ralser, J. Berman, M. Bassilana, R.A. Arkowitz

## Abstract

Treatment failure rates for fungal infections cannot be explained simply by increased rates of drug resistance. Antifungal drug tolerance, the ability of a susceptible isolate to grow in the presence of inhibitory drug concentrations, relies upon a broad set of stress response pathways and can contribute to antifungal drug treatment failures. As the physical properties of the cytoplasm are critical for diverse cellular processes, we investigated whether cytoplasmic mesoscale fluidity is linked to antifungal tolerance, taking advantage of a fluorescent microrheological probe. Here, we show either using fluconazole, a common drug that inhibits ergosterol biosynthesis, or mutants in the ergosterol biosynthesis pathway, that cytoplasmic fluidity decreased and could be reversed upon drug removal. Reducing ribosome concentration decreased drug tolerance and restored cytoplasmic fluidity, highlighting a link between tolerance and cytoplasmic crowding and/or viscosity. However, growth in fluconazole did not increase ribosome concentrations; rather growth in fluconazole increased the number of dormant, hibernating ribosomes, cytoplasmic protein concentration, viscosity and condensate formation. Specifically, growth in fluconazole resulted in a substantial increase in processing bodies (P-bodies), whose presence correlated with azole tolerance. Furthermore, we found a substantial increase in cell-cell heterogeneity in all biophysical, biochemical and molecular outputs analyzed. Together, our results reveal that changes in the physical properties of the cytoplasm occur in response to antifungal drug and suggest that these changes, as well as increased cell-cell heterogeneity, are crucial for survival in fungistatic drugs.

## Introduction

Treatment of fungal infections can fail, as a result of fungal pathogen responses that ultimately permit growth in the presence of antifungal drugs. In general, in *Candida* species, drug resistance levels are relatively low and correlation between clinical outcomes following drug treatment and resistance appears to be lacking^1,2^. This has led to the suggestion that antifungal drug tolerance, which has been observed in a large proportion of clinical isolates and is enhanced at the elevated temperature of the host, explains this discrepancy^2,3^. Drug tolerance refers to the ability of susceptible strains to grow, typically at reduced rates compared to resistant isolates, in drug concentrations above the minimum inhibitory concentration (MIC)^1,2,4^. Tolerance is typically thought to be transient, reversible, non-genetically encoded and generally observed in a subpopulation of varying size^1,2^. Another characteristic of fungal tolerance is that growth in supra-MIC drug levels is reduced compared to growth of resistant strains, possibly reflecting the intrinsic phenotypic heterogeneity of a fungal strain. Recent studies have implicated multiple stress-response pathways and epigenetic processes in antifungal drug tolerance, in particular with respect to the fungistatic azole drugs in the human opportunistic fungal pathogen *Candida albicans*^1–5^.

Despite the increased awareness of the importance of antifungal drug tolerance, with respect to fluconazole (FLC) in *C. albicans*, the cellular changes that occur during survival in this fungistatic drug are largely unknown. Changes in cytoplasmic fluidity have been shown to alter growth rates and cellular metabolism in a range of organisms^6–11^. Cytoplasmic fluidity can be modulated both by macromolecular crowding and viscosity. The bulk of cytoplasm is comprised of particles at the mesoscale (diameters of 10 – 100’s nm) and crowding dominates dynamics at the mesoscale but not at the scale of individual proteins, *i.e.* the molecular or microscale. Viscogens, including polysaccharides, as well as increased protein concentration affect the dynamics of particles across all size scales. Increased macromolecular crowding and viscosity reduces diffusion in cytosol and can perturb a plethora of biological functions^7,12^, in particular reaction rates, protein complex formation and cytoskeletal polymerization rates in different organisms^8,10,13^.

In this study we took advantage of a genetically encoded microrheological probe (GEMs)^6^ to investigate the cytoplasmic mesoscale diffusivity in FLC-tolerant *C. albicans* cells. We show that in these cells there is a striking decrease in cytoplasmic fluidity, which is detectable upon and correlates with ergosterol depletion by antifungal drugs or genetically, and is reversible upon azole removal. This liquid-like to solid-like cytosolic transition is not solely due to increased macromolecular crowding as ribosome levels, visualized and quantified by cryo-EM and mass spectrometry, do not increase in FLC tolerant cells, despite the increase of hibernating ribosomes. However, reduction of macromolecular crowding, either by reducing ribosome levels or by disrupting P-body condensates that form in FLC tolerant cells, restores cytoplasmic fluidity and reduces FLC tolerance. Collectively, these data indicate that reduction of cytoplasmic fluidity and concomitant increase in cytoplasmic heterogeneity in response to ergosterol depletion, *via* fluconazole-dependent inhibition of its target the lanosterol 14-alpha-demethylase Erg11, promotes *C. albicans* growth in high, inhibitory levels of this fungistatic drug.

## Results

### High levels of fluconazole for extended times result in a dramatic decrease in cytoplasmic fluidity

To determine whether exposure to the commonly used antifungal drug, FLC alters cytoplasmic fluidity we took advantage of passive microrheological probes (GEMs)^6^ to measure mesoscale cytoplasmic diffusivity in cells treated with this fungistatic drug. These nanometer-scale particles were used tracers whose movements were followed over time, to measure the local mechanical properties of the cytoplasm. *C. albicans* cells expressing 40-nm GEMs were imaged every 10 msec using HILO microscopy in order to obtain images for single particle tracking and determine the effective diffusion coefficient, D_eff_, which is inversely proportional to viscosity and crowding for Brownian motion. Viscosity and crowding are distinct physical properties that are related, with the former being due to intermolecular interactions whereas the latter results from macromolecules taking up physical space, both of which limit or slow diffusion. False colored temporal projections of GEM trajectories in representative budding cells grown in the liquid media in the presence and absence of FLC are shown in Figure 1A. We used FLC at 0.5X minimum inhibitory concentration (MIC) and 10X-MIC (FLC tolerance conditions) and different incubation times to differentiate between azole-dependent inhibition (0.5X-MIC FLC for 8 Hr) and drug tolerance (10X-MIC for 24 Hr). Figures 1B and S1A show that under conditions of FLC tolerance (high levels of azole), there was a striking decrease in cytoplasmic fluidity (decrease in GEM D_eff_), with an intermediate reduction in GEM D_eff_ after 8 Hr at 10X-MIC FLC and the maximum reduction in GEM D_eff_ was observed after 24 Hr with ∼5-fold decrease (∼ two-fold higher than the background values in fixed cells). As expected, we did not observe substantial cell inviability upon growth in FLC. At the longer incubation times, characteristic morphological changes associated with drug tolerance, *i.e.* enlarged and multi-budded/lobed cells, were observed (Fig. 1C). The decrease in cytoplasmic fluidity was observed irrespective of the promoter used for GEM expression (Fig. S1B). Together, these results suggest that *C. albicans* FLC tolerance is associated with the transition of the cytoplasm from a fluid-like to a more solid-like state.

**Figure 1.**
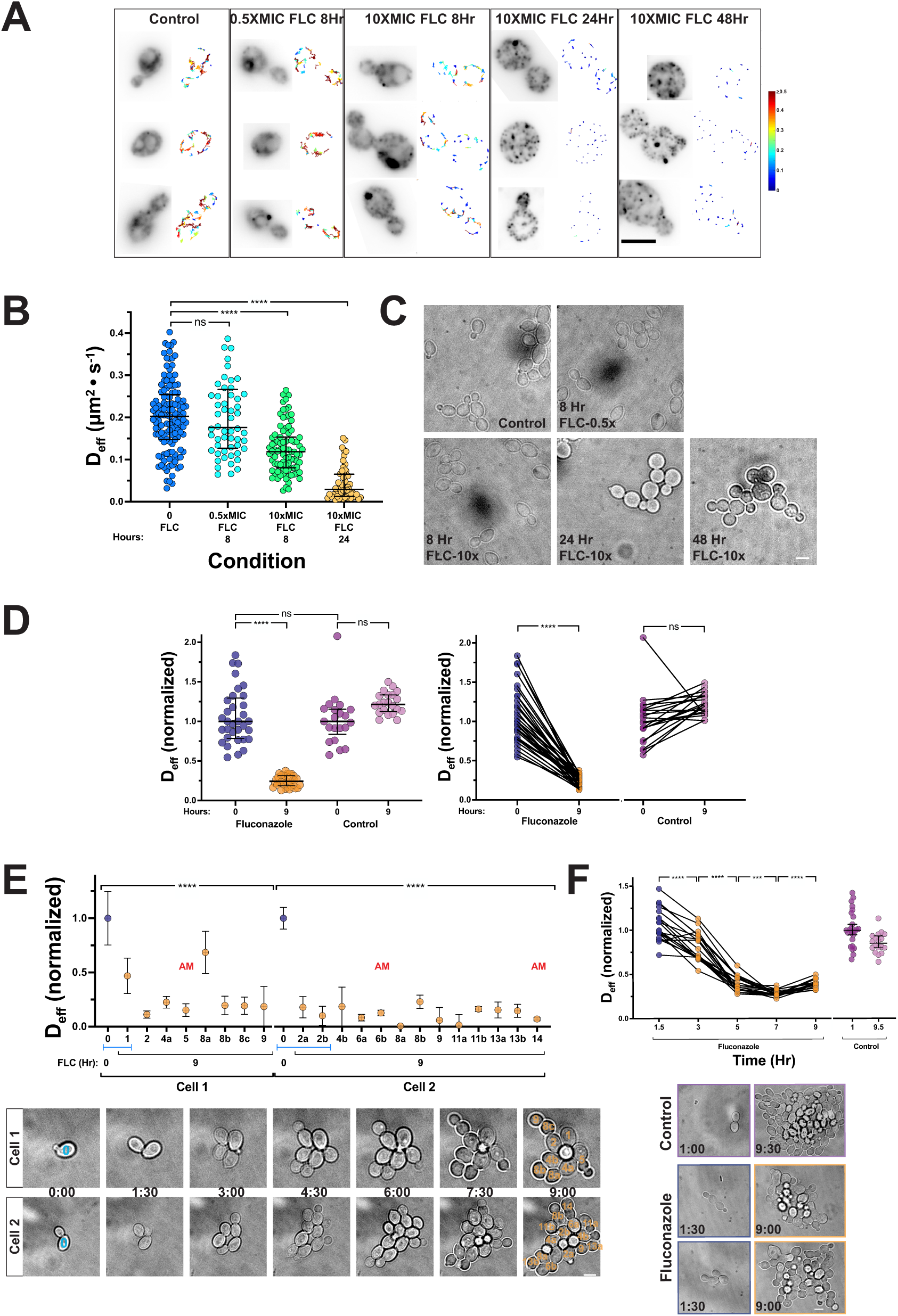
Dramatic decrease in mesoscale cytoplasmic fluidity in cells grown in the presence of high levels of fluconazole that does not correlate with reduced cell growth, morphology alterations or cell age for extended times. A) GEM trajectories. Representative sum projections of GEM acquisitions (left panels) of cells grown in the absence or presence of indicated fluconzole concentrations for indicated times with Deff color coded (right panels) shown. False colored LUT indicating Deff shown. B) Dramatic decrease in GEM Deff grown in high FLC concentration for 24 Hr. Cells were treated as above and each symbol is median cell Deff (*n* = 30 - 125 cells; 15-70 trajectories/cell). Medians and interquartile ranges are indicated; ns not significant, ** < 0.01 and **** < 0.0001. Comparison of cells exposed to FLC for 24 Hr and controls revealed a 1.8-fold increase in the D_eff_ coefficient of variation with FLC. C) Cell morphology is altered during growth in high FLC concentration. Representative cells grown in the indicated conditions. D) Dramatic decrease in mesoscale cytoplasmic fluidity after growth in the presence of FLC for 9 Hr. Cells were grown on agarose pad either in the absence or presence of 10X-MIC FLC and GEM trajectories were determine in the same cells initially and after 9 Hr at 30°C (scatter plot, left and before-after plot, right). Each symbol is median cell D_eff_ (*n* = 22 - 33 cells; 20-200 trajectories/cell; normalized to the median value of 0 Hr cells) with median and interquartile ranges indicated; ns not significant and **** < 0.0001; paired t-test for right panel. Note that at time zero D_eff_ is median of individual cells, whereas at 9 Hr the values are for microcolonies. Comparison cells grown in the presence and absence to FLC for 9 Hr revealed a 2.6-fold increase in the D_eff_ coefficient of variation with FLC. E) Reduction of cytoplasmic mesoscale fluidity does not correlate with cell morphlogy changes, growth rate or cell age. GEM D_eff_ of different cells (median with SEM indicated; normalized to the median value of 0 Hr cells), top panel. Cells with altered morphology (AM) indicated and initial cell at time 0 and same cell at time 9 Hr indicated by blue bracket. Images of cells grown in the presence of FLC (bottom panel). Two cells giving rise each to microcolonies with initial cell indicated (blue 0) and cell number shown in orange, images corresponding two cells quantitated in top panel. F) Decrease in mesoscale cytoplasmic fluidity occurs gradually over 7 Hr and is evident after 3 Hr. D_eff_ of GEMs in individual cells/microcolonies exposed to FLC (or not, controls) for indicated times (top panel) normalized to median of initial values. Each symbol is median cell D_eff_ (*n* = 20 cells; 50-300 trajectories/cell). Paired t-tests with indicated p values (*** < 0.001 and **** < 0.0001). Images of representative cells at indicated times in the absence and presence of FLC (bottom panel).

To follow the response of individual cells, as well as microcolonies, over time in the presence or absence of antifungal drug, we grew cells on synthetic complete agar pads in the presence or absence of 10X-MIC FLC and increased the acquisition time to 30 msec, both of which increased the reproducibility of GEM D_eff_ determinations. Initially, we examined the same cells on agar pads in the presence and absence of this azole, either initially or after 9 Hr. Figure 1D shows that after 9 Hr in the presence of FLC a substantial decrease (4-fold) in cytoplasmic fluidity was observed in all cells, compared to control cells (in which D_eff_ increased somewhat), with a similar decrease was observed irrespective of whether micro-colonies or individual cells were analyzed (Fig. 1D and S1C). In FLC treated samples the D_eff_ from the GEM trajectories of all cells were similar that of the entire microcolony (Fig. S1D), hence further time-lapse analyses focused on microcolonies. We attribute the somewhat stronger effects on GEM diffusion after 8-9 hours in FLC (compare Fig. 1D to 1B) to be due to growth in synthetic complete media in the agar pad (as opposed to rich liquid media) which was necessary for imaging. In the presence of FLC, we observed ∼20% morphologically aberrant cells, typically cells that appeared to have cytokinesis defects, *e.g.* so-called trimers as well as a marked increase in cell doubling time from that of 85-115 min initially to 150-700 min at the end of the experiment. Analysis of the cytoplasmic mesoscale fluidity in individual cells of microcolonies after 9 Hr growth in the presence of FLC did not reveal a correlation between GEM D_eff_ and growth rate, cell morphology or age (Fig. 1E). Nonetheless in supra-MIC fluconazole increased heterogeneity in microcolony growth rates and GEM D_eff_ was observed (see below). Together, these results raise the attractive possibility that fluconazole tolerance is associated with changes in cytoplasmic mesoscale fluidity.

### Decrease in cytoplasmic mesoscale fluidity occurs concomitant with ergosterol depletion and is reversible

We next asked if the decrease in GEM D_eff_ occurred abruptly or gradually over time. First, we found, the decrease in GEM D_eff_ was progressive in cells incubated with FLC (Figure 1F). After 3 Hr in 10X-MIC FLC, the decrease in GEM D_eff_ was statistically significant; a second experiment with shorter incubation times yielded similar results (Fig. S2A). After ∼7 Hr growth in FLC, cytoplasmic mesoscale fluidity reached a minimum. Furthermore, aberrant cell morphologies, cells with broad bud necks and mother-bud pairs that remained connected (failed cytokinesis), were evident (Fig. 1F). We also observed a small, but statistically significant, increase in GEM D_eff_ after longer incubation times, with a 20-40% increase in GEM D_eff_ at 9 – 11 Hr incubation (Fig S2B). Thus, growth in the presence of fluconazole results in a progressive change in the cytoplasmic fluidity, indicative of a cumulative phenomenon.

As fluconazole inhibits the essential lanosterol 14-alpha-demethylase required for ergosterol biosynthesis, does progressive depletion of this sterol result in a decrease in cytoplasmic mesoscale fluidity? Previously, we observed substantial depletion of plasma membrane ergosterol after cells were incubated in FLC - a live-cell reporter for this sterol was no longer detectable after 3 Hr exposure to this antifungal drug^14^, when a decrease in cytoplasmic mesoscale fluidity is observable. Hence, our results are consistent with the idea that reduction in ergosterol levels trigger changes in cytoplasmic mesoscale fluidity.

*ERG11* encodes the FLC target lanosterol 14-alpha-demethylase. We asked if depletion of the Erg11 enzyme also resulted in decreased cytoplasmic mesoscale fluidity. Indeed, cytoplasmic mesoscale fluidity decreased substantially (∼4-fold) when Erg11 protein was depleted by doxycycline (Dox) repression of the pTet promoter driving the sole *ERG11* copy (Fig. 2A). Repression of Erg11 with Dox decreased *ERG11* mRNA, detected by RT-PCR, ∼10-fold (Fig. 2B)^14^. These results were further confirmed with *erg* deletion mutants, specifically an *erg11* deletion mutant kept alive by inactivation of C-5 sterol desaturase Erg3. A significant decrease in GEM D_eff_ was observed in an *erg3Δ*/*Δ* / *erg11Δ*/*Δ* mutant and an *erg3Δ*/*Δ* strain treated with FLC yet was indistinguishable between an *erg3Δ*/*Δ* strain and an isogenic wild-type strain (Fig. S3A). The decrease in GEM D_eff_ in these deletion mutants was less than that observed upon FLC exposure or Erg11 depletion, suggesting that both ergosterol depletion and toxic sterol byproduct accumulation (*e.g.* intermediates eburicol and 14-α-methyl-3,6-diol)^15^ contribute to changes in cytoplasmic fluidity. Nonetheless, deletion of the *ERG11* gene, repression of *ERG11* transcription, and inhibition of Erg11 enzyme with fluconazole all result in a decrease in GEM D_eff_, indicative of a dramatic decrease in cytoplasmic fluidity.

**Figure 2.**
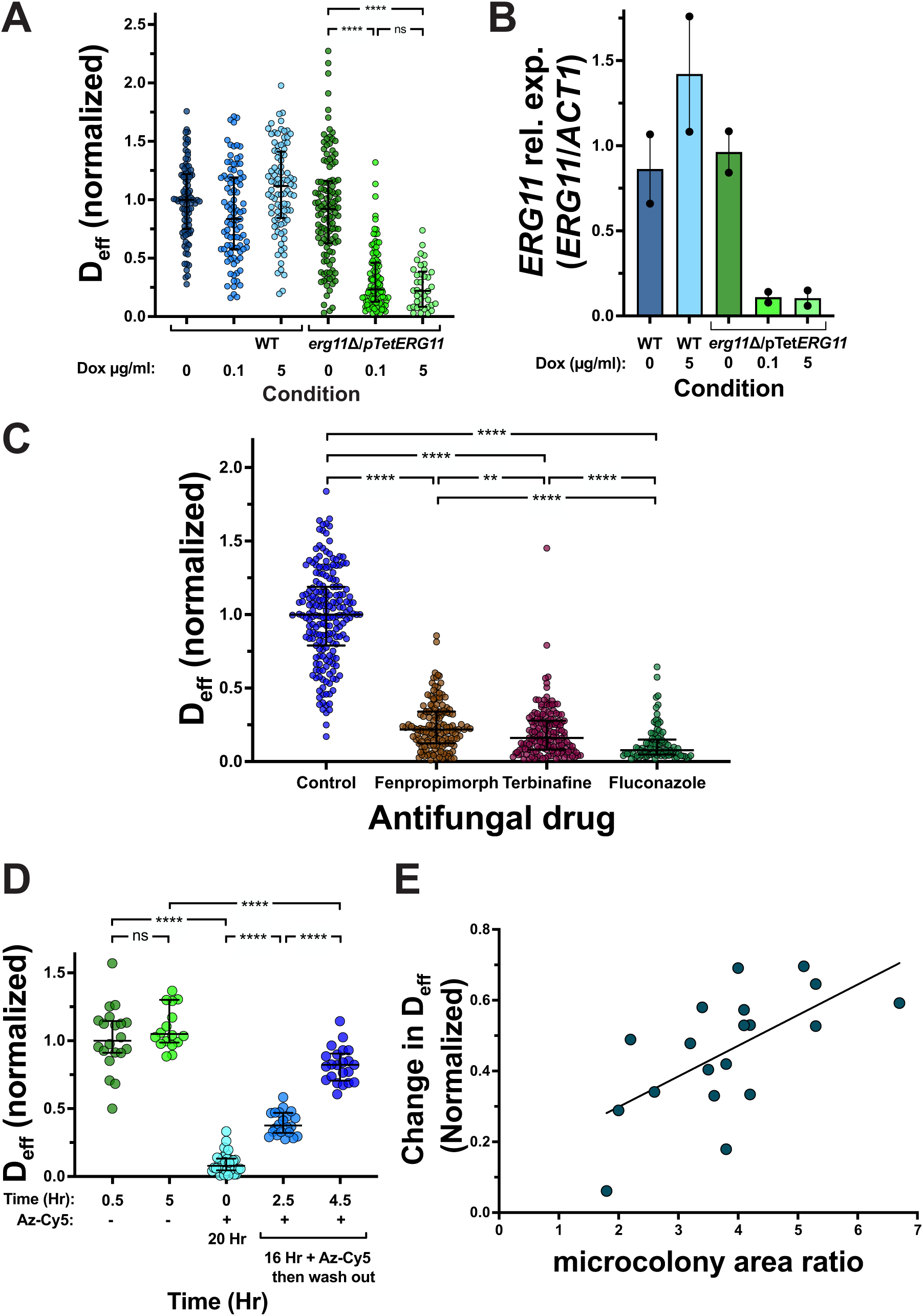
Decrease in cytoplasmic fluidity occurs concomitant with depletion of ergosterol and is reversible upon antifungal drug removal. A) Mesoscale cytoplasmic fluidity decreases upon *ERG11* repression. Cells were grown in the presence or absence of Dox at indicated concentration. Each symbol is median cell D_eff_ (*n* = 40 - 120 cells; 15-450 trajectories/cell; normalized to the median D_eff_) with median and interquartile range indicated; **** < 0.0001. Comparison of cells grown in the presence and absence to Dox revealed a 1.4-1.6-fold increase in the D_eff_ coefficient of variation with FLC. B) *ERG11* expression is reduced upon repression. RT-PCR was carried out using indicated strains (including *erg11Δ/pTetERG11*; PY6786) grown in the presence or absence of Dox, using a CaERG11TM1 primer pair. Values are the means of determinations from 2 independent experiments (points indicate values from each), normalized to *ACT1* and the average level of *ERG11/ACT1* in the absence of Dox was set to 1. C) Inhibition of the ergosterol biosynthesis pathway at different levels all results in a decrease in cytoplasmic mesoscale fluidity. Cells were grown in the presence or absence of indicated antifungal drug, FLC (10 µg/ml), fenpropimorph (1 µg/ml) and terbinafine (10 µg/ml). Each symbol is median cell D_eff_ (*n* = 100 - 200 cells; 15-700 trajectories/cell; normalized to the median D_eff_) with different color shades indicating independent experiments with median and interquartile ranges indicated; ** < 0.01 and **** < 0.0001. Comparison of cells grown in the presence and absence to antifungal drugs revealed a 2- to 3-fold increase in the D_eff_ coefficient of variation with antifungal drug. D) Upon removal of Azole-Cy5 cytoplasmic mesoscale fluidity is restored. Cells were incubated in presence or absence of 1 µg/ml Azole-Cy5 (Az-Cy5) for indicated times and then this antifungal drug removed by centrifugation, washing and incubated on agarose pad further for indicated times. Each symbol is median cell D_eff_ (*n* = 18 - 30 cells; 15-500 trajectories/cell; normalized to untreated median value) with median and interquartile ranges indicated; ns not significant and **** < 0.0001. Note that – refers to absence of Az-Cy5 washout. E) The increase in cytoplasmic mesoscale fluidity upon Az-Cy5 washout is concomitant with growth. The ratio of microcolony area at 4.5 Hr divided by that at 1 Hr was determined and this was plotted with the increase in GEM normalized D_eff_ (D_eff_ at 4.5Hr – D_eff_ at 1 Hr). The linear regression has an r^2^ value of 0.38.

To determine whether the decrease in mesoscale cytoplasmic fluidity was specific to Erg11 function or due to reduced ergosterol in general, we next took advantage of different antifungal drugs, which target distinct steps of the ergosterol biosynthesis pathway. We used the terbinafine (commonly used to treat fungal nail infections), an allylamine to inhibit the squalene epoxidase Erg1 upstream of Erg11, as well as fenpropimorph (an agricultural fungicide), a morpholine which inhibits the two enzymes downstream of Erg1: the C-14 sterol reductase Erg24 and the C-8 sterol isomerase Erg2. Figure 2C and S3B show that these two antifungal drugs also lead to a substantial decrease in cytoplasmic fluidity (∼5-fold for terbinafine and 6-fold for fenpropimorph) without affecting cell inviability. Taken together our results show that a striking decrease in cytoplasmic mesoscale fluidity occurs upon inhibition of the ergosterol biosynthesis pathway, strongly suggesting that ergosterol is critical in this process.

A key characteristic of antifungal drug tolerance is that it is reversible, *i.e*. tolerant cells will grow well in the absence of drug. Hence, we investigated whether the FLC-induced reduction in cytoplasmic mesoscale fluidity is the reversible. To determine the level of this drug, we used a fluorescent azole derivative, azole-Cy5^16^. Cells were exposed to azole-Cy5 at 10X-MIC for 16 Hr which resulted in a similar decrease in GEM D_eff_ as FLC, compared to control cells (Figures 2D and S4A). After removal of azole-Cy5 and incubation in media without drug for 2.5 and 4.5 Hr GEM D_eff_ increased progressively, to >80% of that of the control cells at 4.5 Hr, the time at which azole-Cy5 signal was no longer detectable (Fig. S4B, S4C). Furthermore, we observed a positive correlation (r = 0.62; p < 0.005) between the increase in GEM D_eff_ following drug washout and microcolony growth, indicating that recovery of the wild-type cytoplasmic mesoscale fluidity is associated with cell regrowth (Fig. 2E and S4C). These results reveal that the changes in cytoplasmic mesoscale fluidity are reversible upon removal of drug, which restores ergosterol synthesis and subsequent cell growth.

### Reduction in ribosomes restores cytoplasmic mesoscale fluidity in the presence of fluconazole and reduces antifungal drug tolerance

Ribosomes are one of the major crowders given their abundance in the cytoplasm, as well as the large volume they take up^7,12^, and we recently showed that cytoplasmic mesoscale fluidity during *C. albicans* morphogenesis is regulated by ribosome concentration^11^. Furthermore, rapamycin (RAPA), which reduces FLC tolerance^4^, is also an inhibitor of ribosome biogenesis, *via* the Tor (target of rapamycin) pathway and increases cytoplasmic fluidity in *S. cerevisiae*^6^. To test the hypothesis that there is a link between ribosome levels and antifungal tolerance, we modulated ribosome biogenesis using inhibitors and mutant strains. First, we examined two compounds that inhibit ribosome biogenesis *via* inhibition of transcription of ribosomal proteins and rRNA^17,18^, RAPA and mycophenolic acid (MPA). Tor kinase, inhibits all three nuclear RNA polymerases (Pol I - III), resulting in strong transcriptional inhibition of 35S rRNA precursor, genes encoding ribosomal proteins and 5S rRNA^19^; MPA interferes with rRNA transcription, resulting in the loss of all pre-rRNAs^20^. We used sub-MIC concentrations of RAPA and MPA^21,22^, that did not substantially affect growth or viability. The concentrations of RAPA used have been shown to abolish FLC tolerance but not resistance^4^. RAPA (1 ng/ml) did not alter cytoplasmic mesoscale fluidity, yet at 5 ng/ml it increased fluidity (25% increase in GEM D_eff_) (Fig. 3A, S5A and S5B). Both concentrations of MPA (5 and 30 µg/ml) somewhat reduced GEM D_eff_ by 15% and 25%, respectively (Fig. 3B, S5C and S5D). In cells grown in FLC tolerance conditions (10X-MIC for 24 Hr), both RAPA (1 ng/ml) or MPA (5 µg/ml) restored cytoplasmic fluidity (∼80% of the control GEM D_eff_ with RAPA and ∼30% with MPA) (Fig. 3A, 3B, S5A-D); cytoplasmic fluidity was completely restored with 5 ng/ml RAPA and restored to ∼60% of GEM D_eff_ of that of untreated cells with 30 µg/ml MPA. As RAPA has also been shown to substantially reduce FLC tolerance, measured by the fraction of growth within the zone of inhibition in a FLC disk diffusion assay^4^, we examined whether this was also the case with MPA. Figure 3D shows that MPA similarly results in a reduction of FLC tolerance. Because simultaneous reduction of ribosome concentration in the presence of FLC counteracted the effect of this antifungal drug on cytoplasmic fluidity, we next asked whether reduction of ribosome concentration using RAPA, subsequent to exposure to FLC, would also restore cytoplasmic fluidity. Figure 3C shows that addition of RAPA, after 24 Hr growth in FLC, restored cytoplasmic fluidity ∼60% to that of control after 3 Hr. It should be noted that there was no increase in the percentage of inviable cells when grown with FLC and either RAPA or MPA, respectively. Taken together these results show that immunosuppressive drugs, which reduce ribosome biogenesis, restore normal cytoplasmic mesoscale fluidity in FLC tolerance conditions, and that this restoration is accompanied by a reduction in FLC tolerance, suggesting a link between these two processes.

**Figure 3.**
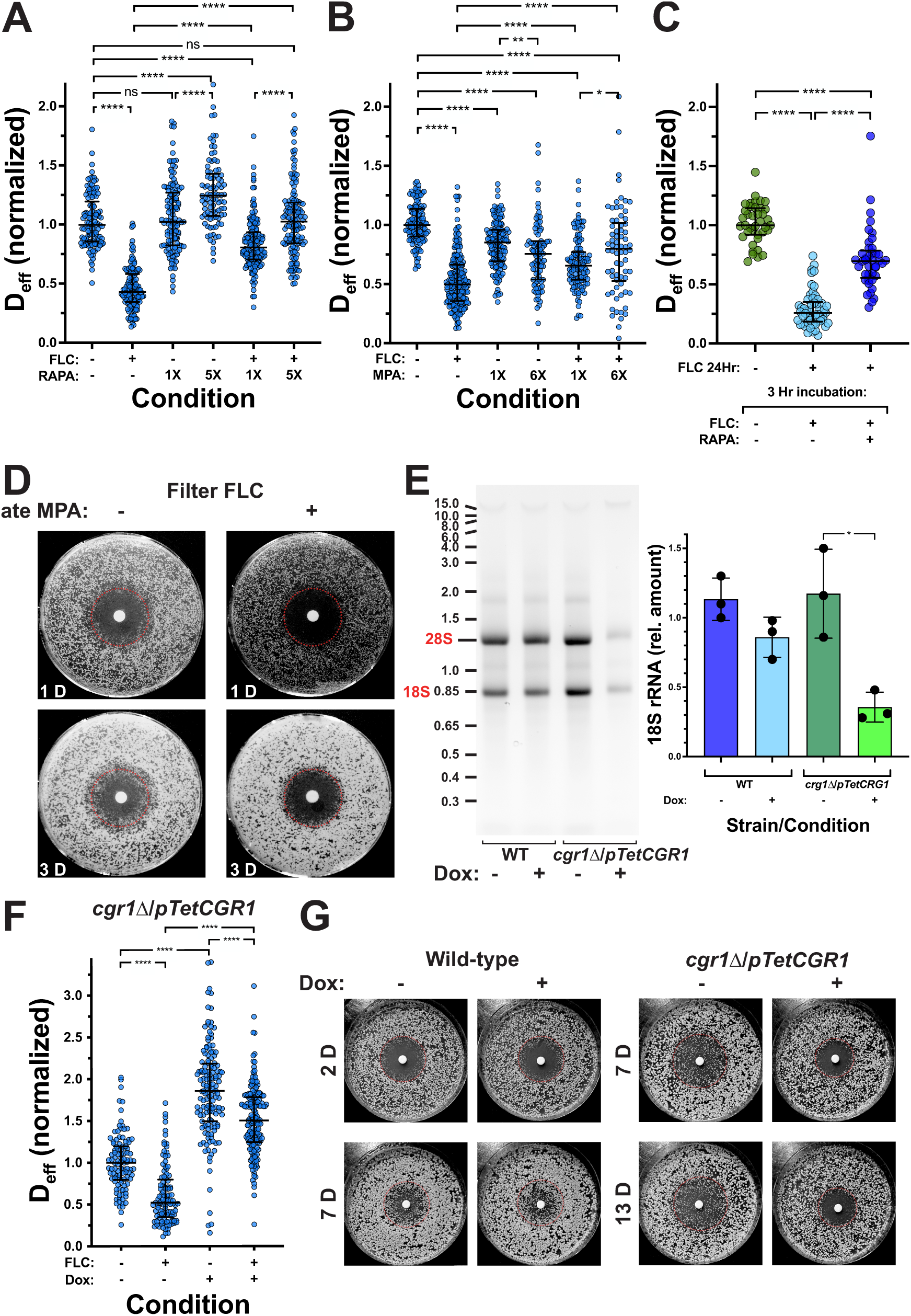
Inhibition of ribosome biogenesis restores cytoplasmic fluidity and reduces fluconazole tolerance. A) Rapamycin restores cytoplasmic fluidity decrease due to FLC. Cells were incubated in FLC and/or rapamycin (RAPA; 1X 1 ng/ml and 5X 5 ng/ml) for 24 Hr. Each symbol is median cell D_eff_ (*n* = 90 - 150 cells; 25-525 trajectories/cell; normalized to median untreated value) with different color shades indicating independent experiments with median and interquartile ranges indicated; ns not significant and **** < 0.0001. B) Addition of rapamycin subsequent to FLC-dependent decrease in cytoplasmic fluidity restores cytoplasmic fluidity. Cells were incubated in presence or absence FLC for indicated time and then rapamycin (5 µg/ml) was or was not added and cells incubated for an additional 3 Hr. Each symbol is median cell D_eff_ (*n* = 40 - 55 cells from 2 independent experiments; 25-400 trajectories/cell; normalized to median untreated value) with median and interquartile range are indicated; **** < 0.0001. C) Mycophenolic acid restores cytoplasmic fluidity decrease due to FLC. Cells were incubated in FLC and/or mycophenolic acid (MPA; 1X 5 µg/ml and 6X 30 µg/ml) for 24 Hr. Each symbol is median cell D_eff_ (*n* = 90 - 180 cells from three independent experiments; 20-740 trajectories/cell; normalized to median untreated value) with median and interquartile ranges indicated; * < 0.05, ** < 0.01 and **** < 0.0001. D) FLC tolerance is reduced by mycophenolic acid. FLC (25 µg) was spotted on filter disks placed on YEPD plate with or without MPA (5 µg/ml) spread with cells and incubated at 30°C for indicated times. The dotted red circle indicates the zone of growth inhibition initially observed and growth within this zone is tolerance. E) Repression of *CGR1* results in a substantial decrease in rRNA. Total RNA was isolated from the indicated strains and analyzed on an agarose gel (left) and quantitation of three independent experiments is shown with mean and SD indicated (right). F) *CGR1* repression overcomes FLC-dependent decrease in cytoplasmic fluidity. Cells were grown in the presence or absence of Dox and FLC. Each symbol is the median cell D_eff_ (*n* = 100 - 160 cells from 2 independent experiments; 15-440 trajectories/cell; normalized to median untreated value) with median and interquartile ranges indicated; **** < 0.0001. G) FLC tolerance is reduced upon *CGR1* repression. FLC (25 µg) was spotted on filter disks placed on YEPD plate with or without 1 µg/ml Dox and spread with cells and incubated at 30°C for indicated times. The dotted red circle indicates the zone of growth inhibition observed after 2D and growth within this zone is tolerance.

As a complementary approach, we used a *CGR1* mutant which affects pre-rRNA processing, reducing 28S and 18S rRNA levels and increases GEM D_eff_^11^. In this *cgr1Δ*/*pTetCGR1* strain the presence of the repressor doxycycline (Dox) resulted in complete repression of *CGR1*, together with a significant decrease in 28S and 18S rRNA levels and an increase in GEM D_eff_^11^ (Fig 3E and 3F). Growth of this strain in FLC tolerance conditions resulted in a substantial reduction in cytoplasmic fluidity that was fully restored upon *CGR1* repression (Fig. 3F). Strikingly, repression of *CGR1* resulted in reduction of FLC tolerance, compared to the wild-type in the presence and absence of Dox, and the *cgr1* mutant in absence of Dox (Fig. 3G). In addition, we observed that the FLC MIC was somewhat decreased in the *cgr1* mutant in the absence of Dox and increased in the presence of Dox, compared to the wild-type strain. Thus, reduction of ribosome biogenesis recovers the decrease in cytoplasmic fluidity in fluconazole exposed cells and reduces antifungal drug tolerance.

### Ribosome levels do not increase in fluconazole tolerant conditions, but there is a significant increase in hibernating ribosomes

Decreasing ribosome levels can restore both cytoplasmic mesoscale fluidity and FLC tolerance, raising the attractive possibility that drug tolerance is mediated *via* regulation of ribosome concentration. Three complementary approaches were used to assess ribosome concentration: liquid-chromatography tandem mass spectrometry (LC-MS/MS) to determine the relative abundance of core ribosomal proteins, total RNA isolation to determine rRNA levels and cryogenic electron microscopy (cryo-EM) coupled with 2D template-matching (2DTM) to quantify 60S ribosomal subunits to directly assess ribosome levels and compositional states^23^. Surprisingly, in contrast to scRNA-Seq results^24^, when cells were grown in 10X-MIC FLC for 24 and 48 Hr we observed little to no change in core ribosomal protein levels (Fig. S6A). However, Figure S6B shows that in FLC tolerance conditions there was a substantial drop in rRNA with only 41% ± 9% (*n* = 4) 18S rRNA remaining. Treatment with RAPA resulted in little detectable rRNA, with 11% ± 10% 18S rRNA, further reduced in cells with both FLC and RAPA (4% ± 7% 18S rRNA). Similar results were observed with MPA, albeit with smaller reductions (Figure S6B). Lastly, quantitation of the cytoplasmic 60S ribosome subunit using 2DTM cryo-EM revealed a small reduction in ribosome levels in FLC tolerance conditions (Fig. 4A - 4C). Furthermore, we analyzed ribosomal compositional states by 3D classification^25^ to determine if there were differences between FLC exposed and control cells (Fig. 4D and 4E). In control cells the majority of ribosomes were in mRNA decoding states (∼40%), whereas, strikingly in FLC-exposed cells the majority of ribosomes were in a hibernating state^26,27^ (∼35%) with a concomitant, almost two-fold decrease in a decoding state (Fig. 4D and 4E). These hibernating ribosomes were defined by the lack of tRNA and the presence of eEF2 and were devoid of a nascent polypeptide, in contrast to decoding, peptidyl transfer and tRNA translocation states (Fig. 4D and S6D). Furthermore the hibernating ribosomes did not appear to be associated with a membrane bound organelle or have a specific location in the cell, as was previously observed in *S. pombe* upon prolonged glucose depletion in which hibernating ribosomes were associated with mitochondria^27^. Closer inspection of these cryo-EM images revealed however that in some cells, ribosomes were found within membranous structures, both within and outside of vacuoles, indicative of autophagy (Fig. 4B and S6C). In addition, we observed repetitive arrays of electron dense material with smallest diameters of ∼100 nm, that were largely devoid of ribosomes (Fig. 4B), which we hypothesize is cellular material that has undergone a phase transition. Together, these results demonstrate that, despite the importance of ribosome levels in FLC tolerance, the concentration of this crowder is not increased during sustained growth in this antifungal drug.

**Figure 4.**
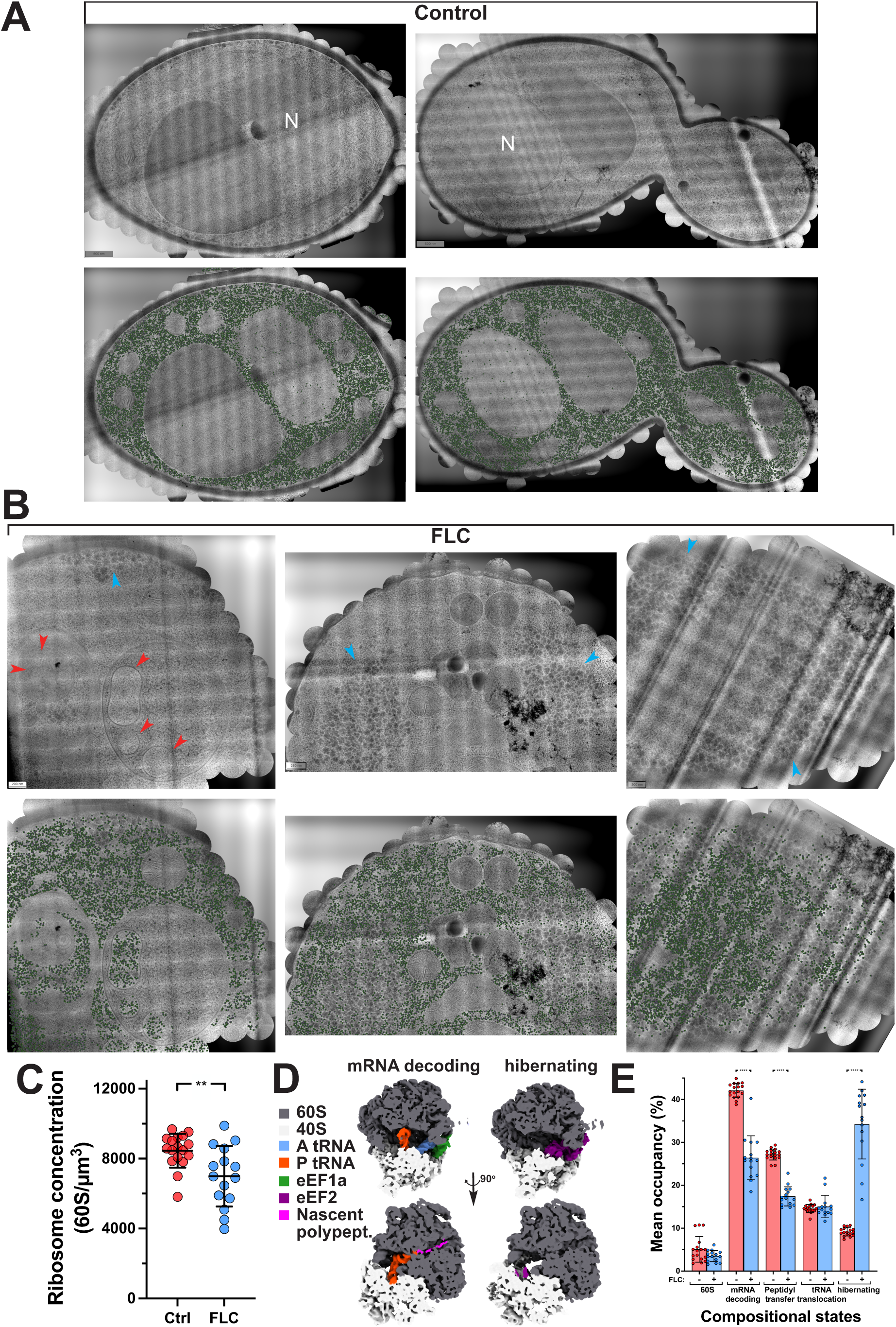
Cryo-electron microscopy shows similar ribosome levels in fluconazole tolerant cells and reveals condensates these cells. A) Montage of cryo-EM exposures of two representative control cells, N is nucleus (upper panels). Overlays of 2DTM detections of the 60S ribosomal subunit (green) in manually segmented cell region (lower panels). B) Montage of cryo-EM exposures of three representative FLC tolerant cells with 60S ribosomal subunit indicated revealing condensates. Top images cryo-EM exposures and bottom images with 60S ribosomal subunit (green) overlays. Red arrow heads indicate regions of autophagy, *i.e.* membrane enclosed structures containing ribosomes and blue arrowheads aggregates. Scale bars indicated. C) Quantitation of ribosome concentration in FLC tolerant cells. Cryo-EM 2DTM was used to determine the concentration of the 60S ribosome (*n* = 15 - 17 cell) with mean and SD shown. D) Ribosome compositional heterogeneity determined by *in-situ* cryo-EM. Two distinct classes of 80S ribosomes display compositional heterogeneity. The top view, isosurface rendering, sliced in front of the tRNA binding sites and the bottom view, isosurface rendering, sliced in front of P-site tRNA and ribosome exit tunnel. Color key for displayed components. E) Ribosome compositional states differ in FLC tolerant cells. Ribosome states determined by *in-situ* cryo-EM and distinct classes corresponding to different stages of translation elongation display compositional heterogeneity. Percentage of different ribosome states. Mean and SD occupancy determined from 17 and 15 FLC-exposed and control cells, respectively; **** < 0.0001. Comparison the occupancy for hibernating ribosome state in cells in presence and absence of FLC revealed a 8.4-fold increase SD with FLC.

### Growth in fluconazole increases cytoplasmic density and viscosity

In addition to being affected by crowders such as ribosomes, cytoplasmic fluidity is inversely proportional to its viscosity, which could also be altered during FLC tolerance. We assessed the intracellular density of cells in FLC tolerance conditions used quantitative phase imaging (QPI), which coupled with cell segmentation, yields surface density (cell dry mass/area). Figure 5A shows that the median cell surface density increased by 30% in FLC – we did not observe a subpopulation of cells, *e.g.* larger or with aberrant morphologies associated with increased surface density. Increased cytoplasmic density is due to changes in water and solute concentrations, which are linked to turgor pressure^28,29^. Hence, we determined whether the turgor pressure was increased in cells grown in FLC tolerance conditions. Figure 5B show that while cell shrinkage in response to increased osmolarity was slightly reduced at all sorbitol concentrations, turgor pressure was indistinguishable from that of untreated cells. Our results suggest that there is an increase in cytoplasmic protein concentration in FLC tolerance conditions and raise the possibility that phase transition, promoted by increased cytoplasmic density, are important during drug tolerance.

**Figure 5.**
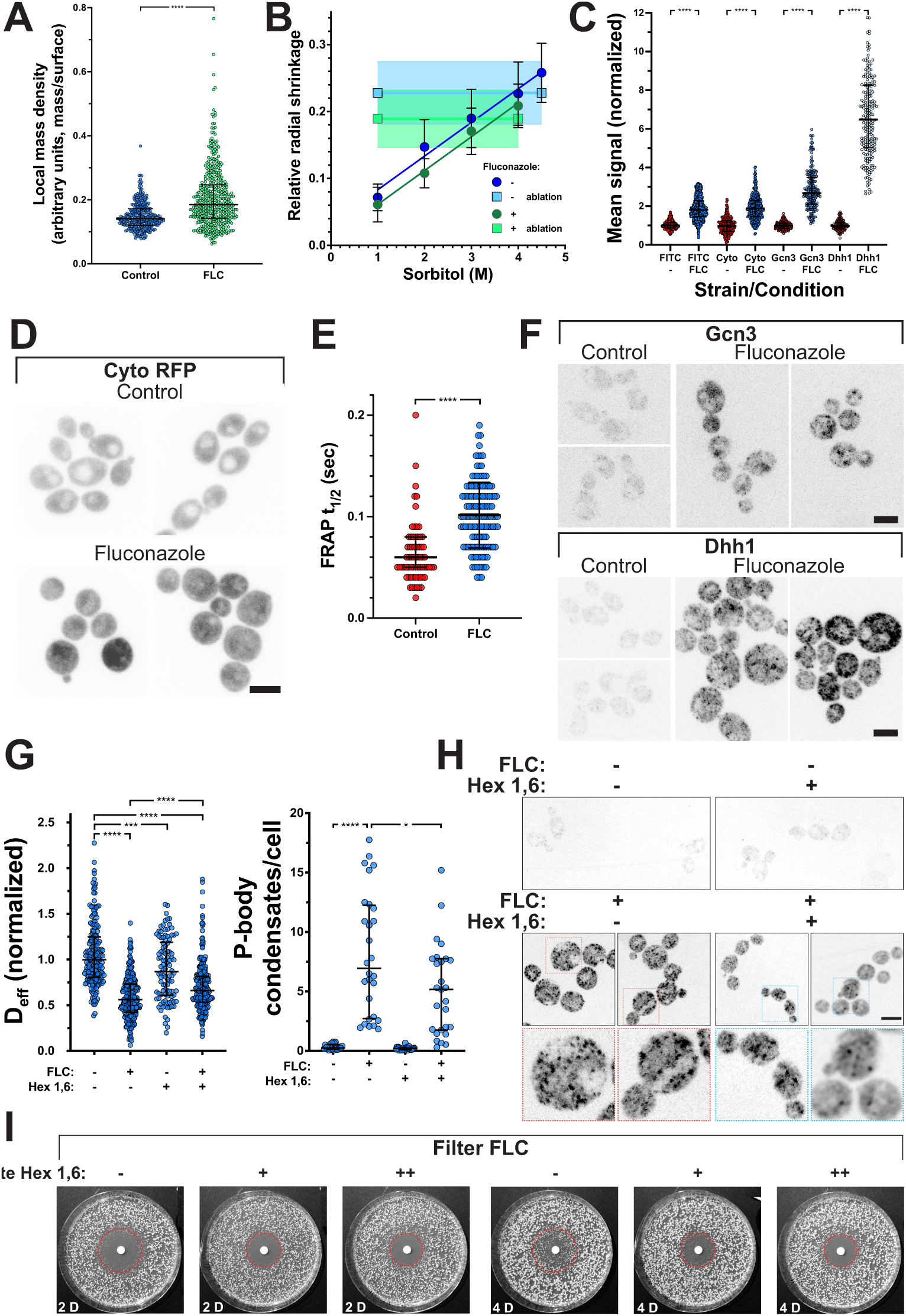
Growth in fluconazole results in a dramatic increase in cytosolic protein concentration together with the formation of processing body condensates that are critical for cytoplasmic fluidity and drug tolerance. A) An increase in surface density occurs upon growth in FLC. Cells were grown in the presence of absence of 10X-MIC FLC for 24 Hr and quantitative phase images were acquired (*n* = 315 - 450 cells from 2 independent experiments). Medians and interquartile ranges indicated; **** < 0.0001. Comparison of the surface density in presence and absence of FLC revealed a 1.6-fold in SD increase with FLC. B) *C. albicans* turgor pressure is unaffected by growth in FLC. Relative radial shrinkage (elastic strain of cell wall) as function of different sorbitol shocks compared to that observed upon cell photo-ablation. The intersection of the two curves is used to estimate the external molarity needed to bring turgor to zero, and thus estimates turgor (*n* = 20 - 40 cells for each sorbitol concentration and *n* = 11 - 18 cells for ablation) with mean and SD indicated. C) Cytosolic protein concentration is dramatically increased in cells grown in FLC. Total protein concentration was determined using FITC staining and cytosolic RFP expression (Cyto) driven by the *ADH1* promoter was also used. Both Gcn3- and Dhh1-3xRFP fusions were used to quantitate their levels, all cells grown in the presence or absence of 10X-MIC FLC for 24 Hr (*n* = 150 - 800 cells; from 3 independent experiments), with medians and interquartile ranges indicated with **** < 0.0001. Comparison of the signal in cells in presence and absence of FLC revealed a 2.7-fold, 1.5-fold and 10.2-fold increase in SD with FLC for FITC labeled proteins, cytosolic RFP and Dhh1, respectively. D) Cytoplasmic RFP levels are increased in cells grown in FLC. Images (sum projections of 6 z-sections) of representative cells grown with or without FLC. E) The mobility of cytosolic GFP is substantially reduced in cells grown in FLC. FRAP of cytosolic GFP in cells (*n* = 70 - 130 cells from two independent experiments). Medians and interquartile ranges indicated with **** < 0.0001. F) Dhh1 P-body condensates are dramatically induced in cells grown in FLC tolerance conditions. Images (sum projections of 6 z-sections) of representative cells expressing Gcn3- or Dhh1-3xRFP grown in the presence or absence of FLC. G) 1,6-hexanediol disruption of P-bodies partially restores mesoscale cytoplasmic diffusivity. Cells were grown in the presence of absence of 10X-MIC FLC for 24 Hr and then treated with or without 5% 1,6-hexanediol (*n* = 120 - 400 cells from 2 independent experiments; 15-240 trajectories/cell; normalized to median untreated value). Medians and interquartile ranges indicated, with ns not significant, **** < 0.0001 and *** < 0.001 (left panel). Right panel, quantitation of the number of P-bodies per cell in the presence and absence of 10X-MIC FLC for 24 Hr and treatment with 1,6-hexanediol. P-bodies were detected using the SPADE algorithm and numbers were extracted per field of view (*n* = 21 - 28 fields of view) with an average of 32 cells per field of view. Comparison of the numbers of P-bodies in presence and absence of FLC revealed a 22.3-fold increase in SD with FLC. H) 1,6-hexanediol disrupts P-body condensates induced by FLC. Images of cells treated identically as above with maximum projections of 6 z-sections shown with enlarged insets (dotted red and blue boxes). I) FLC tolerance is reduced by 1,6-hexanediol. FLC (25 µg) was spotted on filter disks placed on YEPD plate with or without 1,6-hexanediol (0.5 and 1% wt/vol), spread with cells and incubated at 30°C for indicated times. The dotted red circle indicates the zone of growth inhibition initially observed and growth within this zone is tolerance.

To probe cytoplasmic protein concentration in FLC-exposed cells, we examined total cellular protein using FITC staining in fixed cells and cytosolic RFP levels in live cells. The median cellular protein and cytosolic RFP concentration increased two-fold in FLC-exposed cells (Figure 5C and 5D), indicating a substantial increase in cytosolic protein concentration and hence density. As viscosity has been shown to increase with increasing cytoplasmic protein concentration^8,30^, we used fluorescence recovery after photobleaching (FRAP) of cytosolic GFP as a readout for diffusion in the cytoplasm, which is inversely proportional to viscosity^10^. Figure 5E shows that the FRAP t_½_ of cytosolic GFP is increased by approximately two-fold in cells grown in 10X-MIC FLC for 24 Hr. Together, these data indicate that the increase in cytoplasmic density in FLC exposed cells results from an increase protein concentration and we speculate that the associated increased viscosity promotes biomolecular condensation.

### Growth in fluconazole promotes condensate formation

To investigate the effect of FLC tolerance conditions on condensates such as stress granules and P-bodies, we examined Gcn3, the alpha subunit (regulatory subcomplex) of translation initiation factor eIF2B (eIF2Bα); the guanine-nucleotide exchange factor for eIF2, the Pab1-binding protein Pbp1 that regulates TORC1 signaling^31,32^, and the cytoplasmic DEAD-box helicase and mRNA decapping activator Dhh1 that regulates translational repression^33,34^. In *S. cerevisiae*, Gcn3 and Pbp1 localize to stress granules, which form in response to a range of stresses, most notably glucose starvation-induced cytoplasmic acidification (GS-ICA)^35–38^, while Dhh1 is found predominantly in P-bodies, which are involved in degradation and/or storage of mRNAs^39,40^. In *C. albicans* several stresses trigger the formation of Dhh1 condensates, including glucose deprivation and osmotic stress^41^. We generated strains in which an endogenous copy of either of these three genes, *GCN3, PBP1* and *DHH1*, was fused to 3xRFP. For each resulting protein fusion, an increase in signal was observed in FLC treated cells concomitant with a decrease in cytoplasmic mesoscale fluidity; the median cell signal of the Gcn3 and Pbp1 stress granule reporters, increased > two-fold, whereas a substantial 6-fold increase was observed with the P-body reporter Dhh1, (Fig. 5C, 5F, S7A, S7B, S8B). Condensates were observed with Gcn3-3xmSc and to a lesser extent with Pbp1-3xmSc expressing cells in FLC tolerance conditions yet were much more marked in Dhh1 expressing cells, with multiple P-bodies observed in some cells (Fig. 5F and S7A). We also examined whether these P-bodies were dynamic and how FLC affected such dynamics. Given the low signal of the Dhh1 RFP fusions, we acquired images at a single optical plane every 0.6 sec. Despite substantial photobleaching of the signals, we observed apparent movement of Dhh1 P-bodies at these time scales (Fig. S8A). Strikingly, when cells were grown in FLC tolerance conditions, Dhh1 P-bodies were no longer dynamic. Together, these results indicate that in FLC tolerance conditions, there is an increase in cytoplasmic protein concentration and biomolecular condensation, in particular with respect to P-bodies, which suggests increased levels of polysomes due to changes in the ribosome mRNA transit rate^42^.

In *S. cerevisiae,* upon GS-ICA, the cytoplasm undergoes a dramatic transition from a fluid-like to a solid-like phase - the result of increased molecular crowding and condensation of the cytoplasm, including formation of Gcn3 aggregates^38^. In *C. albicans*, following GS-ICA, *i.e.* 2 Hr incubation with 2-deoxyglucose and antimycin A, we observed for each of the three reporters (stress granules Gcn3-3xRFP and Pbp1-3xRFP, and P-bodies Dhh1-3xRFP) an increase in median cell fluorescence signal ranging from 1.5 - 2.4-fold (Fig. S6C, S6D and S8B). For the Gcn3 expressing strains condensates were observed upon GS-ICA, which was not the case with Dhh1 and Pbp1, in which condensates were condensates were somewhat more evident in GS-ICA conditions (Fig. S7C, S7D and S8B). To determine whether the FLC-induced phase transition was also affected by energy depletion, we compared the decrease in cytoplasmic fluidity in FLC tolerance conditions to that of GS-ICA. GS-ICA resulted in ∼ two-fold greater decrease in cytoplasmic mesoscale fluidity, compared to FLC conditions, and combining treatments resulted in a synergistic decrease (Figure S9A). Hence, our findings, consistent with that of *S. cerevisiae*, indicate that GS-ICA increased cytoplasmic molecular crowding and condensation in *C. albicans*, which is more pronounced than from FLC exposure. We also examined the effect of glucose depletion alone, as incubation with 2-deoxyglucose, primarily results in ATP depletion, leading to protein condensation in *S. cerevisiae*^43,44^, yet inhibits the droplet growth of a synthetic phase separation reporter^45^. Figure S9B shows that while incubation in 2-deoxyglucose alone for either 2 or 4 Hr had little to no effect on GEM D_eff_, these conditions partially restored the cytoplasmic fluidity in FLC conditions, consistent with its role in inhibiting condensate growth. Together, our results suggest that the GS-ICA-dependent cytoplasmic transition is distinct from the changes in the cytoplasm observed in FLC tolerance conditions.

To determine whether disrupting FLC-dependent P-bodies alters cytoplasmic mesoscale diffusivity and drug tolerance, we used 1,6-hexanediol, which has been shown to interfere with weak interactions in biomolecular condensates^46,47^. Cells were grown for 24 Hr in the presence or absence of 10X-MIC FLC and then incubated for 10 min with or without 5% 1,6-hexanediol. Treatment with 1,6-hexanediol alone had only a small effect on GEM mesoscale diffusivity, however addition to cells exposed to FLC for 24 Hr partially restored GEM mobility (Fig. 5G). This increase in mesoscale diffusivity was accompanied by a reduction in the number of P-body condensates per cell (Fig. 5G, 5H). Strikingly, the presence of 1,6-hexanediol (0.5 and 1%) also reduced FLC tolerance (Fig. 5I), suggesting a link between P-body condensate formation, reduced cytoplasmic mesoscale fluidity and drug tolerance. We did not observe an increase in inviable cells grown overnight in 1,6-hexanediol (0.5% and 1%) and FLC compared to FLC alone, indicating that this compound was not increasing FLC-dependent cell death. Our data indicate that reducing cytoplasmic fluidity by altering crowding either by changing ribosome or P-body condensates levels reduces FLC tolerance. Together, our results reveal that tolerance to the antifungal drug FLC is in part mediated by changes in cytoplasmic fluidity to a more solid-like phase, as a result of increased protein concentration and biomolecular condensation.

### Cell-cell heterogeneity in fluconazole-exposed cells

FLC tolerance is characterized by microcolonies that appear slowly at supra-MIC, and those that divide a few times, yet do not form visible colonies^4^. Consistent with the idea that there are subpopulations of cells that respond differently to the drug, two distinguishable transcription patterns were observed after 2 days of FLC exposure^24^. Here, we asked if cells at supra-MIC FLC concentrations exhibit a higher level of cell-to-cell heterogeneity of cytoplasmic properties at the mesoscale (GEM D_eff_, ribosomal states, and condensates) and at the molecular or microscale (individual proteins). Overall, in FLC, we observed a ∼ two-fold increase in the heterogeneity (coefficient of variation) in GEM D_eff_ between individual cells and microcolonies relative to no drug controls (Fig. 1B, 1D and S1C). Similarly, the coefficient of variation for GEM D_eff_ was higher when ergosterol was reduced by *ERG11* repression or with other drugs (Fig. 2A and 2C). In addition, the heterogeneity of hibernating ribosome levels was strikingly higher in FLC than in no drug controls (Fig. 4D). Furthermore, in FLC relative to no drug controls, there was a similar two-fold increase in the heterogeneity of cytoplasmic contents, including: cell density (cell dry mass/cell area), overall protein concentration, cytosolic RFP concentration, and stress granule signals (Fig. 5A and 5C). Specifically, the heterogeneity of P-bodies, detected as signal level or number of Dhh1 foci, increased >10-fold and >22-fold, respectively, in FLC relative to no drug controls (Fig. 5C, and S8B). The increased heterogenetiy of P-bodies did not appear to be a general response to stress, as as only a small increase in hetergeneity (∼2-fold) was observed with glucose stress (GS-ICA; Fig. S8B). Together, these results establish that cell-to-cell cytoplasmic biophysical heterogeneity is a hallmark FLC tolerance at both the mesoscale and microscale.

## Discussion

Antifungal drug treatment failure is frequently due to drug tolerance and resistance, which contribute to infection recurrence. Antifungal drug tolerance is distinct from drug resistance as, for example in the case of fungistatic azoles, tolerance manifests itself as heterogenous growth above the MIC that is thought to be a non-genetically encoded, physiological response^1,2^. Antifungal drug tolerance is defined as a growth phenomenon, and we have little insight into the molecular basis or processes that underly this critical cause of infection treatment failure. Here we show that under conditions where tolerant growth to azoles is observed, *i.e.* extended growth at supra-MIC FLC levels, there is a decrease in cytoplasmic mesoscale diffusivity in the human fungal pathogen *Candida albicans* (Fig. 6). This dramatic change in cytoplasmic physical properties occurs upon ergosterol depletion and is reversible when azole is removed, indicating that this change coincides with tolerance. Growth in supra-MIC FLC levels results in an increase in cytosolic protein levels, together with a significant increase in P-bodies and a corresponding increase in cytosolic viscosity (Fig. 6). Intriguingly, FLC tolerance is decreased, and cytoplasmic fluidity is partially restored by reducing macromolecular crowding, either by decreasing ribosome levels, or by disrupting P-body condensates that form in cells grown in FLC, using 1,6-hexanediol. Together, these results suggest that reduction of cytoplasmic fluidity in response to ergosterol depletion, *via* azole-mediated Erg11 inhibition, enables *C. albicans* to proliferate in high levels of fungistatic drug.

**Figure 6.**
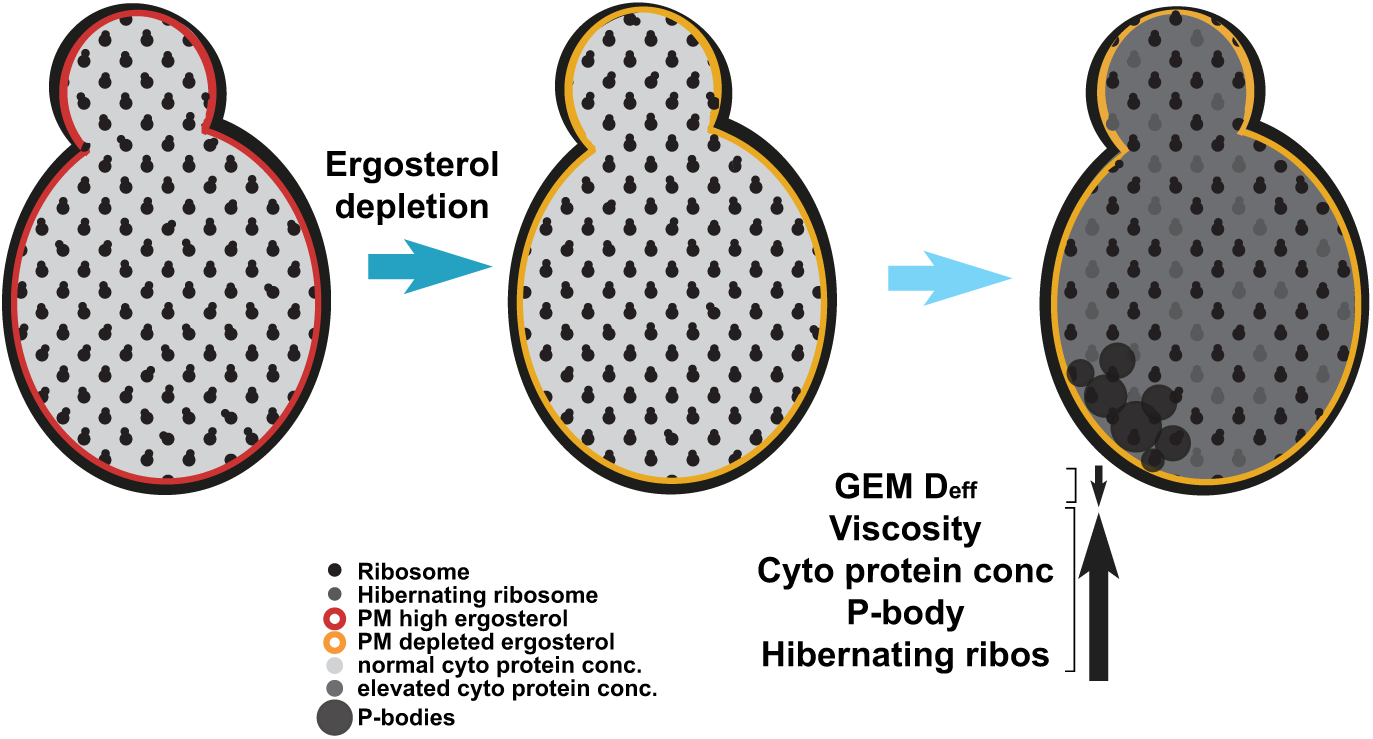
Schematic of cytoplasmic changes that occur in fluconazole tolerant cells. Colors indicate levels of ergosterol in the plasma membrane (PM). Ergosterol depletion is either by inhibition by antifungal drugs or repression of a key enzyme in the ergosterol biosynthesis pathway (Erg11). Different cytoplasmic components are indicated in the key.

Chemical inhibition or genetic repression of Erg11 results in a liquid-like to solid-like cytosolic transition. The decrease in cytoplasmic mesoscale diffusivity is observed concomitant with the reduction of ergosterol at plasma membrane growth sites which occurs over several hours growth. In addition, inhibition both upstream (using terbinafine) and downstream (using fenpropimorph) of Erg11 results in similar changes in the cytoplasm, strongly indicating that it is the depletion of ergosterol that triggers these changes. What could be sensing these changes in cellular ergosterol levels? Studies in *S. cerevisiae* have identified two conserved kinases, the AGC (protein kinase A, G and C) kinase Ypk1 and the CaM (calmodulin) kinase Rck1, which stoichiometrically bind ergosterol^48^. Ergosterol was also shown to stimulate *S. cerevisiae* Ypk1 kinase activity *in vitro*^48^, however ergosterol does not appear to be an essential Ypk1 activator upon sphingolipid depletion^49^. In addition, the level of the Ste11 group Ssk2 homolog, Ssk22, protein depends on ergosterol^48^. These three kinases are candidates for ergosterol sensors critical for changes in cytoplasmic fluidity and, indeed, Ypk1 kinase activity is required for basal FLC tolerance in *Cryptococcus neoformans*^50^.

Notably, tolerant growth is reduced by inhibition of the TOR or calcineurin pathway^4^. It is tempting to speculate that Ypk1, which is a TOR complex-dependent protein kinase, may ultimately link ergosterol levels to changes in cytosolic protein concentration, indeed, this pathway is intimately associated with modulation of both gene transcription and translation, *via* the translation machinery as well as translation initiation^51^. Interestingly, both azole tolerance^4^ and the effects of FLC/ergosterol depletion on the *C. albicans* cytoplasm are reversible, suggesting that these effects are not genetically encoded.

Our data reveal that manipulation of cytoplasmic fluidity, by decreasing the amount of a macromolecular crowder, *e.g.* ribosomes or ribonucleoprotein granules such as P-bodies, reduces tolerance to FLC, suggesting a causal role for cytoplasmic fluidity in tolerance. Single cell RNASeq revealed that after several days in MIC-level FLC, when a small increase in tolerance was observed, a subpopulation of cells with increased expression of mRNA encoding ribosomal proteins and rRNA was observed^24^. However, in substantially higher FLC concentrations (10X-MIC for 24 Hr), we observed reduced rRNA levels (28S and 18S) and similar levels of core ribosomal proteins, when normalized by the median protein level, by mass spectrometry. Furthermore, cryo-EM analyses revealed similar ribosome (60S) levels in cells treated with FLC compared to untreated controls, although there was a 3-fold increase in hibernating ribosomes. Strikingly, ribosomes were detected within membrane bound organelles in cells grown in 10X-MIC FLC, most likely vacuoles and/or multi-vesicular bodies (MVBs)^52^, which indicates substantial autophagy in these conditions. Such autophagy is consistent with the reduction in rRNA we observed yet, given the levels of core ribosome proteins were not substantially reduced in the presence of FLC, this suggests that increased ribosome protein synthesis may balance or counter the ribosome protein degradation by autophagy.

It has been proposed that cytoplasmic macromolecule concentration is tuned to maximize the rates of a range of biochemical reactions^53^. Indeed, such cytoplasmic protein homeostasis has been demonstrated in *Xenopus* egg extracts, in which protein synthesis rates are optimal in the standard cytoplasmic concentration, whereas protein degradation continues to increase as cytoplasmic concentration increases^30^. These observations suggest that the increase in cytosolic protein concentration and viscosity we observed in cells grown in 10X-MIC FLC could potentially result in a ∼ 3-fold decrease in mRNA translation rate, together with ∼1.5-fold increase in protein degradation rate (based upon values determined in *X. laevis* egg extracts^30^). We speculate that growth in 10X-MIC FLC could alter protein translation rates, perhaps *via* the TOR pathway, thereby increasing cytoplasmic protein concentrations initially. In light of this observation of increased protein concentration in the cytosol of FLC tolerant cells, it is notable that the major molecular chaperone Hsp90 is critical for azole tolerance^1,2,4^, suggesting that protein homeostasis plays an important role during this process. However, at the high cytosolic protein concentrations observed in 10X-MIC FLC exposed cells, we anticipate decreased rates of protein synthesis that could result in reduced growth rates and thereby increase chances of survival in this fungistatic drug. Indeed, the increase in inactive, hibernating ribosomes devoid of tRNA and nascent polypeptide, together with the small decrease in overall ribosome concentration are consistent with a decrease in translation. The hibernating ribosome state is likely to preserve the translation machine until cells overcome FLC-induced stress. Phase separation of biomolecular condensates is driven by increases in cytoplasmic protein concentration^54^, hence the increase P-body condensates observed upon prolonged 10X-MIC FLC exposure is likely promoted by the substantial increase in cytosolic protein concentration and corresponding increase in viscosity. The response to FLC, *via* ergosterol depletion, does not appear to be solely a stress response, as we did not observe a similar large increase in two proteins that localize to stress granules, but rather a specific increase in P-bodies. P-bodies are cytoplasmic ribonucleoprotein granules that store and degrade translationally repressed/non-translating mRNAs, as well as proteins involved in mRNA processing and turnover^55,56^. The release of mRNAs from polysomes can result in an increase in P-bodies^42,57,58^, although it appears that reversible storage of non-translating mRNAs in P-bodies is limited and not a general phenomenon with respect to most mRNAs^59^. Glucose depletion leads to polysome collapse and the release of mRNA in *S. cerevisiae*^59^ and to transiently (within 15 min) fluidize the cytoplasm^44^, however at longer times cytoplasmic fluidity decreased. Interestingly, we observed that glucose depletion with 2-deoxyglucose for either 2 or 4 Hr partially restored cytoplasmic mesoscale diffusivity in cells exposed to 10X-MIC FLC, which we attribute to polysome collapse in these conditions. Dissolution of P-bodies formed in cells treated with FLC partially restored cytoplasmic mesoscale diffusivity and reduced FLC tolerance, demonstrating that tolerance is intimately associated with cytoplasmic fluidity. We speculate that decreased cytoplasmic fluidity leads to reduced translation rates, and that tuning cytoplasmic fluidity may be a general means to provide optimal conditions that promote slow growth and antifungal tolerance.

Fluconazole tolerance is characterized by heterogeneity with respect to cell growth and transcriptional patterns^4,24^. Here we show that FLC tolerant cells exhibit a striking cell-to-cell heterogeneity in the cytoplasm, both at the mesoscale (GEM D_eff,_ ribosomes states and P-body condensates) and the microscale (cell density and cytosolic protein concentration) (Fig. 6). It is tempting to speculate that cytoplasmic heterogeneities in crowding and viscosity constitute a bet-hedging mechanism that promotes survival in a cell sub-population exposed to fungistatic drug. Further drug-induced increases in cytoplasmic heterogeneity at the mesoscale and microscale would then diversify sub-populations that can initially survive, providing a positive feedback loop that increases long-term survival and recurrent infections.

## Online Methods

### Strains and media

Strains used in this study are listed in Table S1a. For transformation, strains were grown in YEPD (yeast extract, peptone, dextrose) supplemented with Uridine (80 µg/ml) at 30°C. Cells were grown in liquid YEPD medium, supplemented with Uridine at 30°C and for time lapse microscopy on agarose pads with in synthetic complete (SC) media at 30°C. For doxycycline (Dox) gene repression, YEPD was supplemented with 5 µg/ml Dox. Antifungal drugs, fluconazole, fenpropimorph, terbinafine and Azole-Cy5 (synthesized by WuXi Biologics) stocks were 1 mg/ml (H_2_O), 0.5 mg/ml (DMSO), 5 mg/ml (DMSO), 50 µg/ml (EtOH), respectively. For drug inhibition of ribosome biogenesis, cells were grown with indicated concentrations of rapamycin (5 µg/ml stock in DMSO) or mycophenolic acid (5 mg/ml stock in EtOH), with or without FLC for 24 Hr prior to imaging. For 1,6 hexanediol incubation cells were incubated for 10 min at 30°C, at indicated concentrations using a 10-fold concentrated stock solution. For glucose starvation-induced cytoplasmic acidification (GS-ICA) and glucose depletion, logarithmically growing cells were washed twice with SC lacking a carbon source and resuspended in an equal volume of SC with either 20 mM 2-deoxyglucose or 20 mM 2-deoxyglucose and 10 µM Antimycin A (10 mM stock in EtOH) for GS-ICA. For glucose depletion (only 2-deoxyglucose) cells were incubated for 2 or 4 Hr and for GS-ICA cells were incubated for an additional 2 Hr. Agar pads for imaging contained the same concentration of drug or repressor as overnight incubation. For experiments with sorbitol, budding cells were incubated with the indicated final concentrations of sorbitol prior to imaging.

The oligonucleotides used in this study are listed in Table S1b. Plasmid templates for GEM integration behind *ADH1* or *TEF1* promoter, pFA-CaPfV-GFPγ^A206K^-CdHIS1 and pFA-CaPfV-GFPγ^A206K^-ARG4 were as described^11^. The tetracycline repressible strains (*erg11Δ/pTetERG11* and *cgr1*Δ/pTet*CGR1*) were constructed as described^14^. Strains expressing plasma membrane targeted GFP (GFP-CtRac1) were constructed as described^60^ and strains expressing cytosolic GFP and mScarlet were constructed as described^60–62^. Mutants (*erg3* and *erg11*) in SC5314 background were generated using strain in which both copies of *ARG4* were removed by CRISPR and subsequently the CaPfV-GFPγ^A206K^ construct was integrated. The *ARG4* and *ERG3* deletions were performed using the CRISPR method with the primers, Donor Arg4-FW/Donor Arg4-RV and Donor Erg3-FW/Donor Erg3-RV^63^. The subsequent deletion of *ERG11* was done in the *erg3Δ/Δ* mutant by transient CRISPR with the following modifications: the CAS9-encoding plasmid, pADH147^63^, was modified to encode CAS9 without the selection markers. The forward primer for amplification of the A fragment was replaced by primer, pSNR52-FW and the reverse primer for amplification of the B fragment was replaced by primer, gRNA conserved-RV. For *ERG11* deletion, the 90 bp donor DNA in the original method^63^ was replaced with a ∼5 kb fragment containing the *NAT* selection marker and the maltose-inducible FLP system amplified from plasmid pRB895^64^. Primers Erg11-NAT-FW/Erg11-NAT-RV each contained 58 bp of homology to the 5’and 3’ ends of the *ERG11* ORF, respectively. Gcn3, Dhh1, and Pbp1 3xmSc fusion strains were generated by integration of PCR product using corresponding oligonucleotides and pFA-3xmSc-ARG4.

### Microscopy, sample preparation and image analysis

#### Light microscopy

The GEM nanoparticles were imaged, using TIRF on a Nikon Ti eclipse inverted microscope (Nikon France S.A.S., Champigny-sur-Marne, France) equipped with an iLas^2^ scan head (Roper scientific, Evry, France), an iXon 888 EMCCD camera (Andor technology, Belfast, UK) and a 100x CFI-APO-TIRF oil NA 1.49 objective as described with 30 msec image acquisition and readout time for all experiments except those shown in Figure 1B which used 10 msec image acquisition and readout time. Temperature was controlled with an Okolab incubator (Ottaviano, Italy) at 30°C for all acquisitions. For GEM acquisitions together with spinning disk imaging in the Az-Cy5 experiments the spinning-disk confocal modality on the above-described microscope, equipped with a Yokogawa CSU-W1 (Yokogawa Electric Corporation, Tokyo, Japan), was used with a 642 nm diode laser. Spinning-disk confocal microscopy was carried out described previously^14,61^ with a PLANAPO total internal reflection fluorescence (TIRF) 1.45-numerical aperture (NA) 100 × objective. FRAP experiments were carried as described using a Zeiss LSM 980 confocal microscope with a 63X NA1.4 oil immersion objective with 30 mW diode 405 nm and 488 nm LASERs^11^. Turgor pressure determination was carried essentially as described^65^ on a Zeiss LSM 780 NLO confocal microscope with a Spectra-Physics Mai Tai DeepSee multi-photon laser using PDMS flow through chambers. Multi-positions were acquired with a motorized XYZ stage. Quantitative phase imaging (QPI) was carried out with a SID4Bio (Phasics, Saint Aubin, FR) on a Zeiss Axiovert 200M inverted microscope equipped with a motorized stage and a homemade enclosure for temperature control. Acquisitions were carried out at 30°C on an agar pad using QCAM software (version 2.0.13) and data was analyzed with Phasics software. The effective diffusion of the GEM particles was calculated as described^6^, using the plugin Mosaic^66^ in Fiji (version 1.54f) in a Windows 10/Intel computer and Matlab (version R2023a). Matlab was also used to generate the plots of all trajectories using a colormap. Each track with ≥11 consecutive frames was also drawn in the raw image coordinate frame and color-coded by its D_eff_ on a fixed scale (0–0.5 µm² s⁻¹) using a custom script (scripts available upon request).

Total cellular protein concentration was deterimined using FITC staining essentially as described^11^. Cells were fixed with 70% EtOH for 1 Hr, following washing were stained with 50 ng/mL FITC fo 30 min and subsequently washed. Quantitation of protein concentration using FITC labeling, cytosolic RFP concentration and Gcn3/Dhh1/Pbp1 RFP fusion concentrations all from sum projections was carried out using a custom MATLAB program: individual cells were fitted with ellipses, the mean fluorescence intensity inside each cell was measured, and the background level was subtracted. The background signals were estimated as the average intensity of the darkest 20% of pixels in each frame (scripts available upon request). Numbers of P-bodies were detected and quantified using the SPADE algorithm (https://gitlab.inria.fr/ncedilni/spade) as described^67^. FRAP data was analyzed using easyFRAP, an open-source online FRAP analysis program^68^. Measurements with a mobile fraction < 0.7 and single exponential curve fits with r^2^ values < 0.4 were not analyzed. Cell inviability was determined using propidium iodide staining as described (Vernay et al 2012). Scale bar is 5 µm unless indicated otherwise.

#### Electron microscopy

Cryoplunging was carried out essentially as described^11^. Au grids (200 mesh) with a 2/2 silicone oxide support film were glow-discharged on both sides for 45 sec at 20 mA. Cells suspensions (3 µl; 0.2 OD_600_) were applied to the support film side of grids, blotted for 10 sec and then frozen in liquid ethane using a cryoplunger (GP2 Leica, Wetzlar, Germany) and frozen. FIB-milling was carried out using an Aquilos 2 FIB/SEM (Thermo Fisher, Waltham, MA) with a stage cooled to < -190°C in a 35° AutoGrid sample holder. Grids were sputter-coated with metallic Pt and then coated with organo-Pt essentially as described^23^. An overview of the grid was created by montaging SEM images and isolated cells or cell clusters at the center of grid squares were selected for FIB-milling. Lamellae were generated automatically using the AutoTEM software (Thermo Fisher, Waltham, MA), with the following protocol: rough milling 1: 1 nA; medium milling 2: 0.3 nA (1.0° overtilt); fine milling 0.1 nA (0.5° overtilt); finer milling 0.1 nA (0.5° overtilt); lamella polishing: 50 and 30 pA, 0.2°overtilt resulting in 150 nm thick lamellae which were subsequently sputter-coated with Pt for 5 sec at a current of 5mA.

FIB-milled grids were imaged in a Titan Krios TEM (Thermo Fisher, Waltham, MA) operated at 300 keV and equipped with a BioQuantum energy filter (Gatan, Pleasanton, CA) and K3 camera (Gatan, Pleasanton, CA). The instrument was controlled using SerialEM^69^. Individual lamellae were manually centered in the microscope and then moved to a position 60 µm below the eucentric height to achieve fringe-free illumination. The stage was tiled 15° to compensate for the milling angle and overview images were obtained with a pixel size of 76.8Å. Individual cells were annotated in these overview images using napari^70^ and high-resolution montages were obtained for these cells using DeCo-LACE acquisition scripts^71^. The physical pixel size in high-resolution exposures was 1.05Å, defocus was maintained at 1 µm, and the total exposure was 30 e/Å^2^. The exposures were dose-fractionated into 30 frames.

Movies were imported into the *cis*TEM software package^72^ and motion-corrected using a custom version of unblur^73^ as described^71^, and binned to a final pixel size of 2.0 Å. CTF parameters and sample thickness were estimated using CTFFIND5^74^. The structure of the vacant *C. albicans* 80S ribosome^75^ (PDB code: 7PZY) was modified by deleting subunits corresponding to the 40S subunit and the resulting 60S structure was converted to a density map at 2.0 Å pixel size using the simulate program in *cis*TEM^76^ using a B-factor scaling of 2. 2D template matching of individual exposures was performed using the GPU-accelerated version of the match_template program in the *cis*TEM suite^77^. Rotation angles were search using a 2 in-plane and 3 out-of-plane step-size and defocus values were searched within a 240 nm slab at a 40 nm step size.

The number of 60S ribosome subunits per imaged area was determined by manually segmenting cytosolic areas from the montaged cryo-EM images (Fig. 4A, 4B, and S6C) and dividing the number of detections within the segmentation by the segmented area. The imaged volume was calculated by fitting the thickness of individual exposures, as estimated by CTFFIND5, to a 2-dimensional cubic B-spline model with 3 knots^19^ and integrating the estimated thickness at every pixel of the segmented area.

To quantify the proportions of ribosomes in different states of translations we extracted particle images around 60S detections. Particles from FLC treated cells (n= 123,369) and control cells (n= 167,380) were pooled into a single particle stack. Orientations and positions were refined by a single round of “Manual Refinement” in *cis*TEM. The particle stack was binned to a pixel size of 8.0 Å and particle parameters were exported into the Frealign format. Frealign (v 9.11_Aug2017)^78^ was then used for classification of the particle stacks into 20 classes, without refinement of particle position or orientation, for 100 iterations. A spherical focus mask with a radius of 80 Å was placed over the A-site/GTPase (center coordinates in Å: 216.7,259.4,347). The resulting 20 reconstructions were compared to states resulting from in-situ single particle analysis in *S. cerevisiae*^25^ and based on this grouped into “60S”, “mRNA decoding”, “Peptidyl transfer”, and “tRNA translocation”. We observed classes that did not fit into these groups and which we instead designated as “Hibernating”, based on the presence of eEF2 and absence of tRNA^26,27^. The best fitting structure for this class in PDB is 8K2D without eIF5a^79^. We then summed the mean occupancies of particles from FLC treated or control cells for all classes belonging to each group. For visualization, we recalculated reconstructions of representative classes from particles at a pixel size of 2.0 Å using Frealign and filtered the reconstructions to a resolution of 7.0 Å.

### Proteomics

Sample preparation was essentially as described^11,80^. Quality control samples to monitor LC-MS performance were created from pooling small aliquots of all samples. Peptide quantities were estimated *via* Quantitative Fluorometric Peptide Assay (Pierce). LC-MS based proteomic data acquisition was performed as described^81^. In brief, samples were injected on a ACQUITY M-Class HPLC (Waters) connected to a ZenoTOF 7600 mass spectrometer with an Optiflow source (SCIEX), separated on a HSS T3 column (300 µm×150 mm, 1.8 µm; Waters) using a 20 min active gradient. We used a Zeno SWATH acquisition scheme with 85 variable-sized windows and 11 msec accumulation time. LC-MS raw data was processed using DIA-NN 1.8^82^. First, a spectral library was predicted including the UniProt Proteome of *C. albicans* SC5314 (UP000000559), as well as the sequence of the genetically encoded multimeric nanoparticle (see description above). For the main search, we enabled tryptic digestion allowing for one missed cleavage, no variable modification, N-terminal Methionine excision and carbamidomethylation as fixed modification of Cysteines. Mass accuracies were set to fall within 20 ppm and match-between-runs was enabled with protein inference on Protein level. The obtained report was processed using Python 3.9 with the pandas (1.4.3) and NumPy (1.23.0) packages. Data was filtered to less or equal than 1% FDR concerning Global.Q.Value, as well as PG.Q.Value and Lib.PG.Q.Value. Prior to plotting, protein group quantities were sample-wise median normalized in two steps: first, subtracting the sample median in log2-space, then subtracting the log2-sample median of entities not belonging to the ribosome, according to UniProt annotated protein names (*i.e.* containing “60S” or “40S, that is the 76 core ribosome proteins).

### Disk diffusion assays

Disk diffusion assays were carried out essentially as described^4^ and carried out on YEPD agar plates containing uridine. A sterile circular filter disk was placed in the center of each plate and 25 µg of FLC spotted on the filter disk. Plates contained no drug, MPA, Dox or 1,6-hexanediol and were imaged after incubation at 30°C for the indicated time.

### RNA extraction and RT-PCR

Cells were grown in YEPD media in the presence or absence of different compounds. RNA extraction and RT-PCR were carried out as described^83^. Oligonucleotide pairs ACT1pTM/ACT1mTM and ERG11pTM/ERG11mTM, respectively.

### Statistical analysis

Data were compared by the Mann-Whitney U test and where relevant the paired or unpaired t-test using GraphPad Prism (v. 8) software, with all *p* values indicated in figure legends. Unless stated otherwise medians and interquartile ranges are indicated. Pearson correlation coefficient and simple linear regression were determined using GraphPad Prism (v. 8) software.

### Data availability

All data supporting the findings of this study are available within the paper and its Supplementary Information. Cryo-EM micrographs will be deposited into the EMPIAR database and mass spectrometry data into PRIDE database, released upon publication.

## Acknowledgements

We thank H. Labbaoui and S. Bogliolo for assistance, the PRISM Imaging facility (B. Monterroso and S. Ben-Aicha) and the Microscopy Imaging Cytometry d’Azur (MICA) for microscopy support (the Zeiss LSM980 was acquired with funds from Conseil départemental - Alpes-Maritimes CD06 and Inserm ITMO Cancer), M. Rigney for support with cryo-EM sample preparation, P. Pognonec for use of Phasics Sid4bio. Cryo-EM data were acquired at the UMass Chan Medical School Cryo-EM Core Facility. This work was supported by the CNRS, INSERM, Université Côte d’Azur, ANR (ANR-19-CE13-0004-01), EC (MSCA-ITN-2015-675407; MSCA-IF-2020-101029870; ERC-SyG-2020 951475), the German Ministry of Education and Research (BMBF) MSCoreSys (031L0220), FRM (SPF202309017657 and FDT202404018460) and NIH (TL1TR001454) grants.

## Notes

### Competing Interest Statement

The authors have declared no competing interest.

