## Supplementary Figures 1-9 and Supplementary Table 1 for "Tolerance to the antifungal drug fluconazole is mediated by tuning cytoplasmic fluidity"

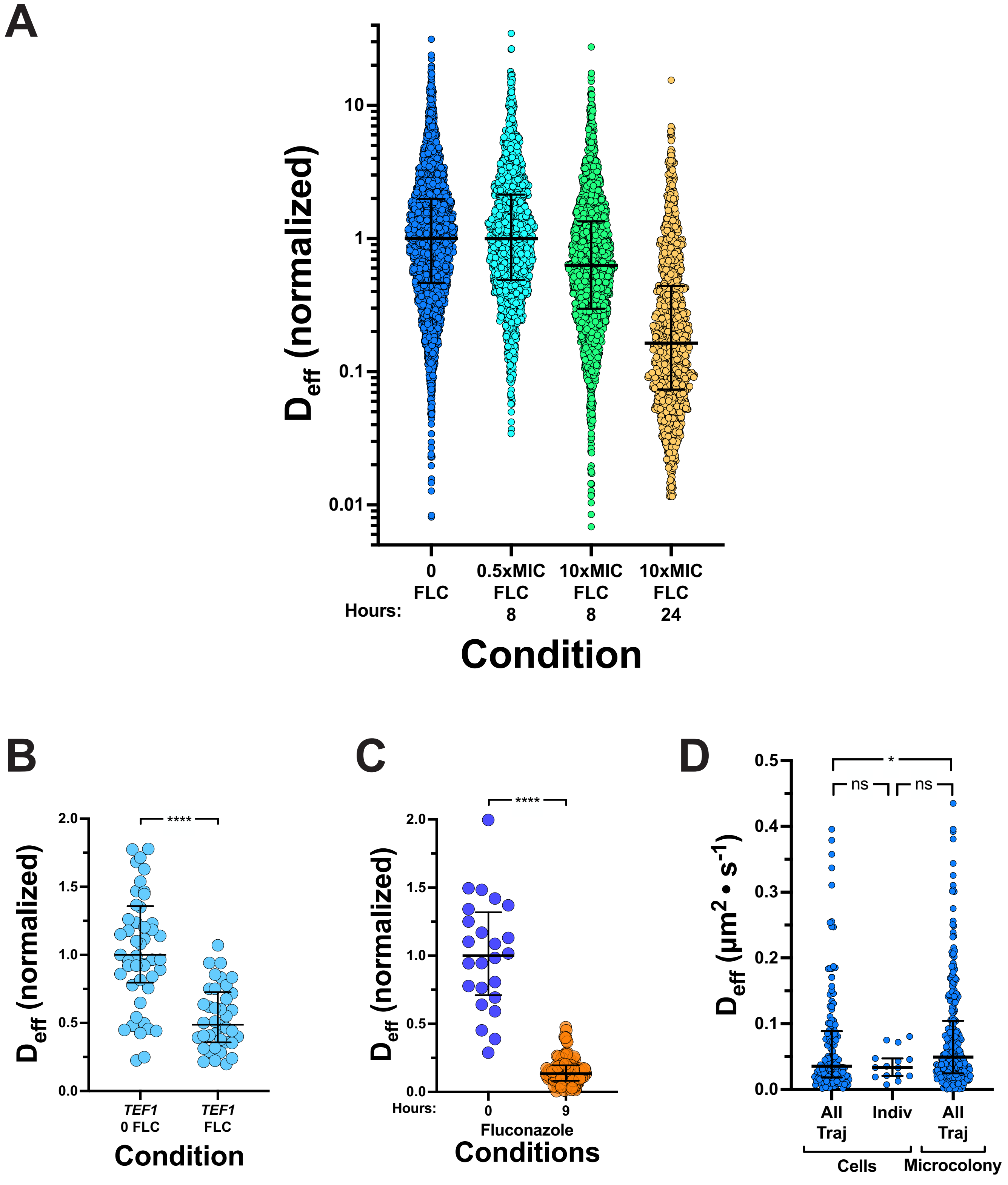


**Figure S1. The effect of fluconazole on GEM effective diffusion is independent of expression level similar in single cells and microcolonies.** A) All trajectories of cell exposed to FLC. Effective diffusion of all trajectories from cells in the absence or the presence of FLC for indicated times. Cells from Fig. 1B, with 740 - 4050 trajectories for each condition. Medians and interquartile ranges are indicated. B) The effect of FLC is GEM effective diffusion is independent on their expression level. Each symbol represents the median $D_{eff}$ of indicated cells ($n$ = 38-49 cells each condition; 20 -180 trajectories per cell), expressing GEMs under the control of the *TEF1* promoter incubated with or without FLC for 24 Hr, with medians and interquartile range indicated (normalized to time 0); **** < 0.0001. C) Analysis of the effect 10X-MIC FLC on individual cell GEM $D_{eff}$. GEM $D_{eff}$ of individual cells from one experiment in Fig. 1D are shown with medians and interquartile ranges are indicated (normalized to initial time); **** < 0.0001. Comparison of $D_{eff}$ coefficient of variation from individual cells grown in the presence and absence to FLC for 9 Hr revealed a 1.5-fold increase with FLC. D) Analyses of individual cropped microcolony cells or entire microcolony yields similar GEM effective diffusion. Analyses of cropped cells, either cell $D_{eff}$ medians ($n$ = 14 cells; 3 – 19 trajectories per cell) or all trajectories and microcolony cluster. Medians and interquartile ranges are indicated, with * < 0.05 and ns not significant.

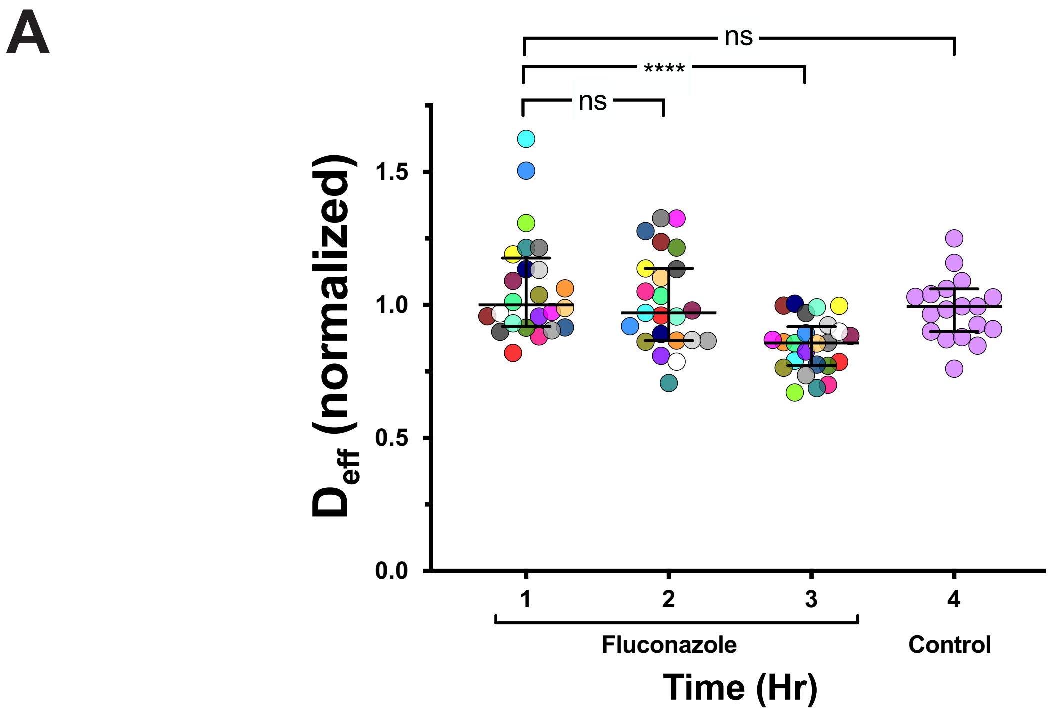


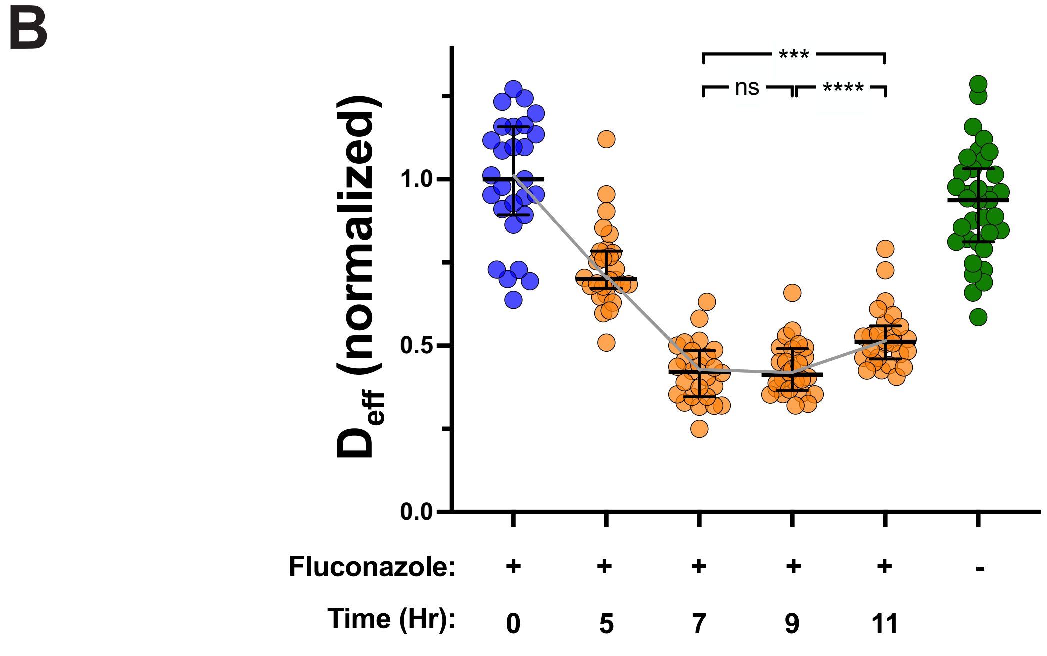


**Figure S2. Cytoplasmic mesoscale diffusivity decrease is evident after 3 hours in the presence of fluconazole begins to increase after 9-11 hours.** A) Decrease in cytoplasmic mesoscale fluidity is observed after 3 Hr in FLC. $D_{eff}$ of GEMs in cells/microcolonies exposed to FLC (or not, controls) for indicated times as in Fig. 1F. Each symbol is median cell $D_{eff}$ normalized to initial time (*n* = 19-24 cells; 40 - 1100 trajectories/cell) with interquartile ranges indicated. Paired t-tests with indicated p values (ns not significant and **** < 0.0001 for FLC treated samples). B) Decrease in mesoscale cytoplasmic fluidity is followed by a small increase in fluidity after 9-11 Hr. Deff of GEMs in cells/microcolonies exposed to FLC (or not, -) for indicated times as in Fig. 1F. Each symbol is median cell $D_{eff}$ normalized to initial time 0 (*n* = 26-35 cells; 20 - 1050 trajectories/cell) with interquartile ranges indicated. Paired t-tests with indicated p values (ns not significant, *** < 0.001 and **** < 0.0001).

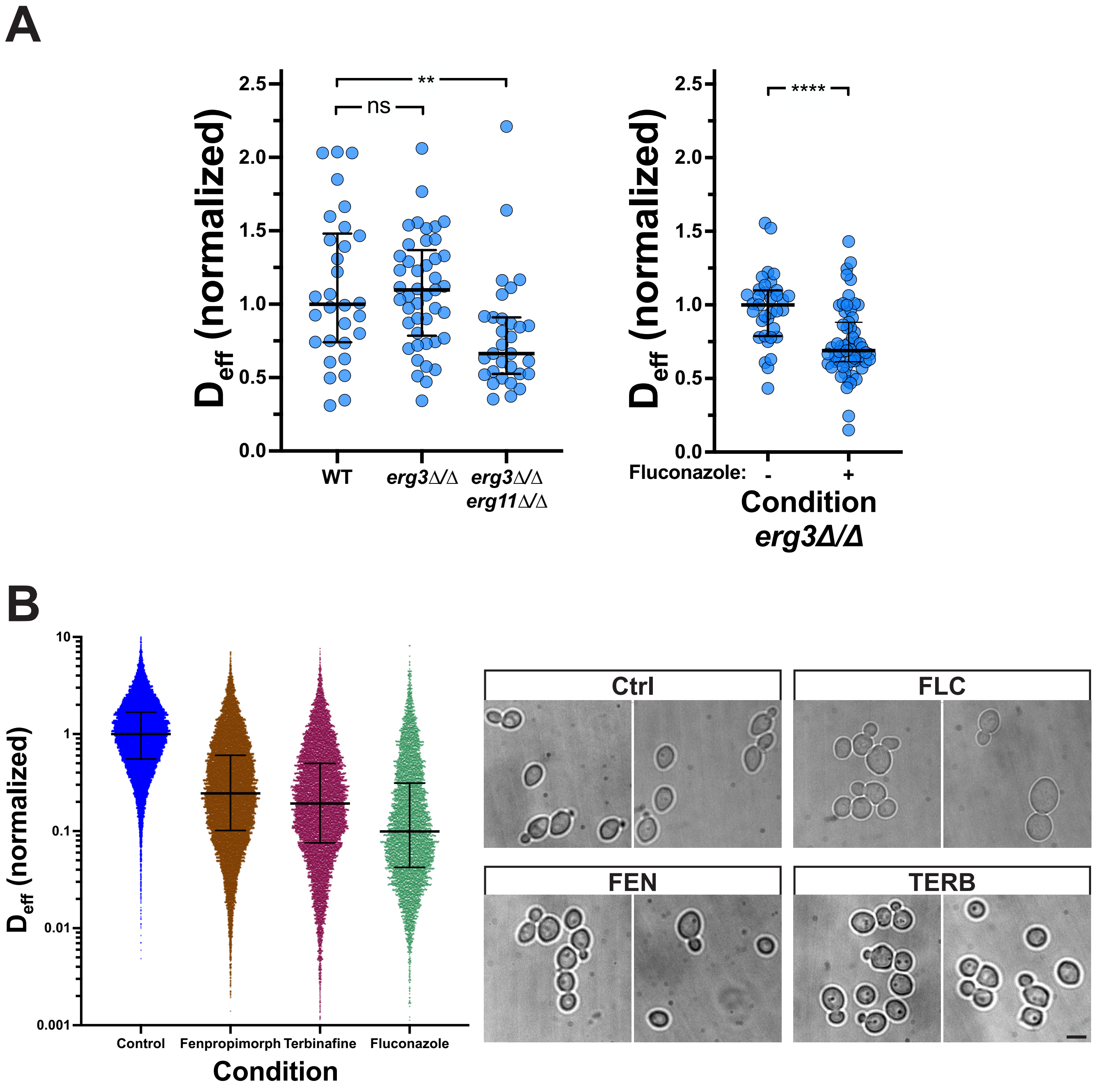


**Figure S3. Ergosterol depletion as well as removal of *ERG11* result in a decrease in cytoplasmic fluidity.** A) Deletion of *ERG11* results in a decrease in cytoplasmic fluidity. Indicated strains were grown in the presence or absence of FLC. Each symbol is median cell $D_{eff}$ normalized to wild-type or control cells ($n$ = 30-59 cells; 10 - 260 trajectories/cell) with interquartile ranges indicated; **** < 0.0001. B) All trajectories of cell exposed to fenpropimorph, terbinafine and FLC. Effective diffusion of all trajectories from cells in the absence or the presence of indicated antifungal drugs (normalized to control cells). Cells from Fig. 2C, with 6425 - 22100 trajectories for each condition (right) and images of representative cells shown (left). Medians and interquartile ranges are indicated; **** < 0.0001.

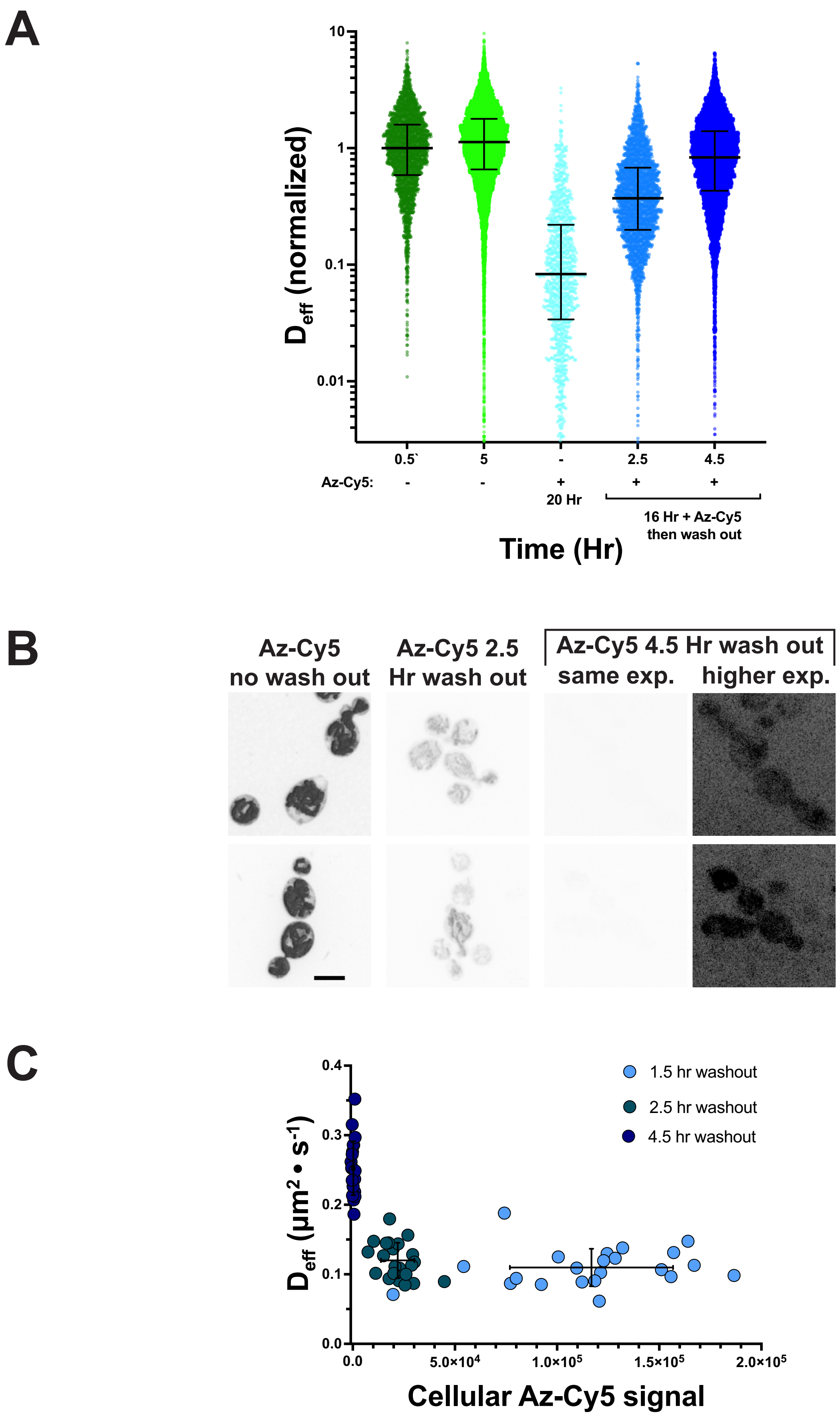


**Figure S4. Decrease in cytoplasmic fluidity is reversible upon azole removal and cell growth.** A) All trajectories of cell exposed to Azole-Cy5 followed by drug removal. Effective diffusion of all trajectories from cells in the absence or the presence of indicated Az-Cy5 for indicated times (normalized to untreated cells). Cells from Fig 2D, with 1050 - 11320 trajectories for each condition. Medians and interquartile ranges are indicated. B) Cellular azole levels are rapidly reduced following drug removal. Images of cells incubated with Az-Cy5 for 16 Hr and then incubated an additional (no wash out) or with drug removed and incubated for an additional 2.5 and 4.5 Hr. The farthest right images shown at higher exposure to reveal cells. C) GEM $D_{eff}$ as a function of cellular Azole-Cy5 levels. Mean cellular Az-Cy5 signal were determined from sum projections (26 z-sections) along with the cell's median GEM $D_{eff}$. Bars indicated SD and mean.

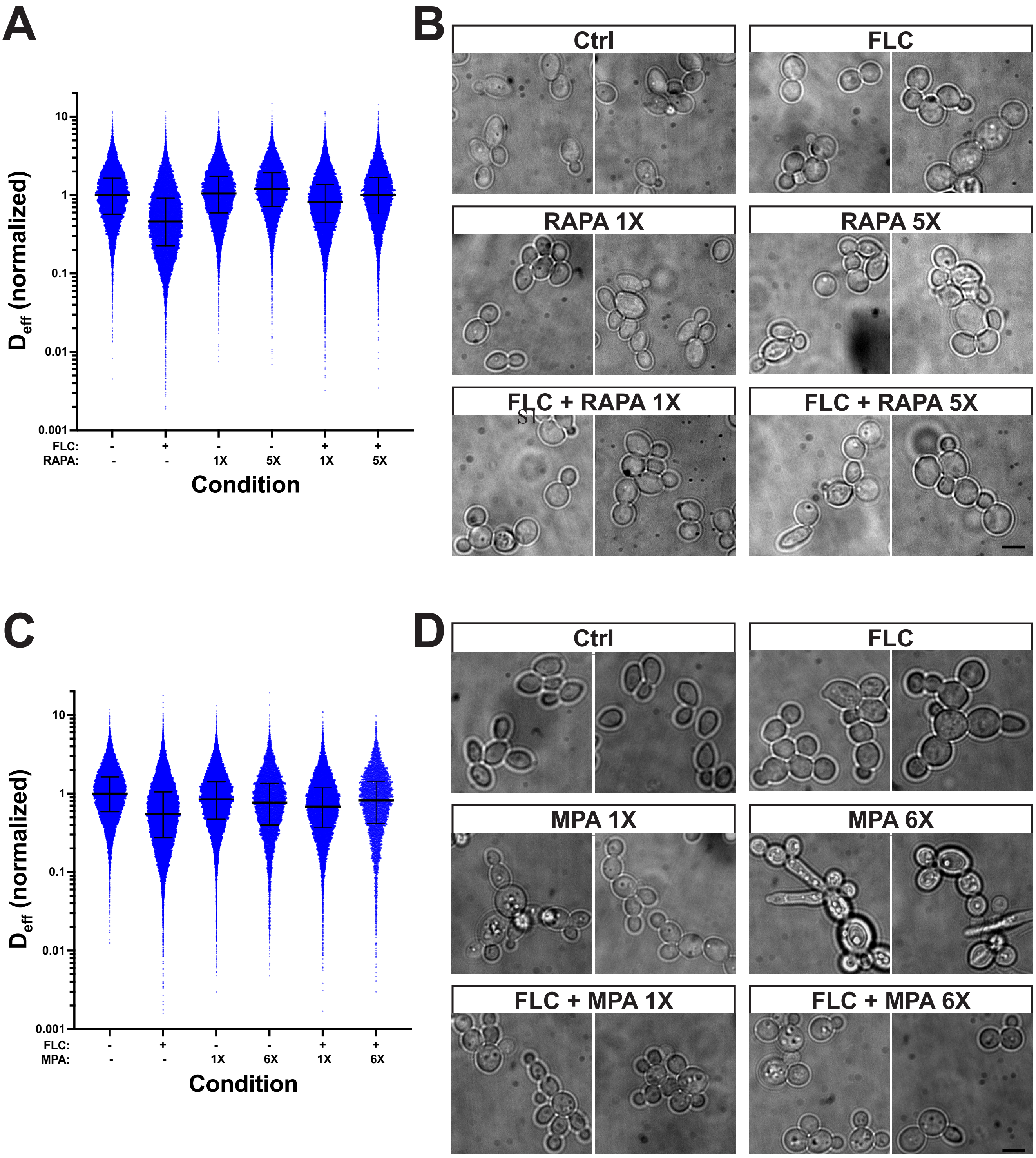


**Figure S5. Ribosome biogenesis inhibitors restore fluconazole induced decrease in cytoplasmic fluidity.** A) All trajectories of cell grown in rapamycin with and without FLC. Effective diffusion of all trajectories from cells in indicated conditions (normalized to untreated cells). Cells from experiment in Fig. 3A with 14470 - 19530 trajectories for each condition. Medians and interquartile ranges are shown. B) Morphology of cells grown in FLC and/or rapamycin. Images of cells from experiment in Fig. 3A. C) All trajectories of cell grown in mycophenolic acid with and without FLC. Effective diffusion of all trajectories from cells in indicated conditions (normalized to untreated cells). Cells from experiment in Fig. 3B with 6170 - 25180 trajectories for each condition. Medians and interquartile ranges are shown. D) Morphology of cells grown in FLC and/or mycophenolic acid. Images of cells from experiment in Fig. 3B.

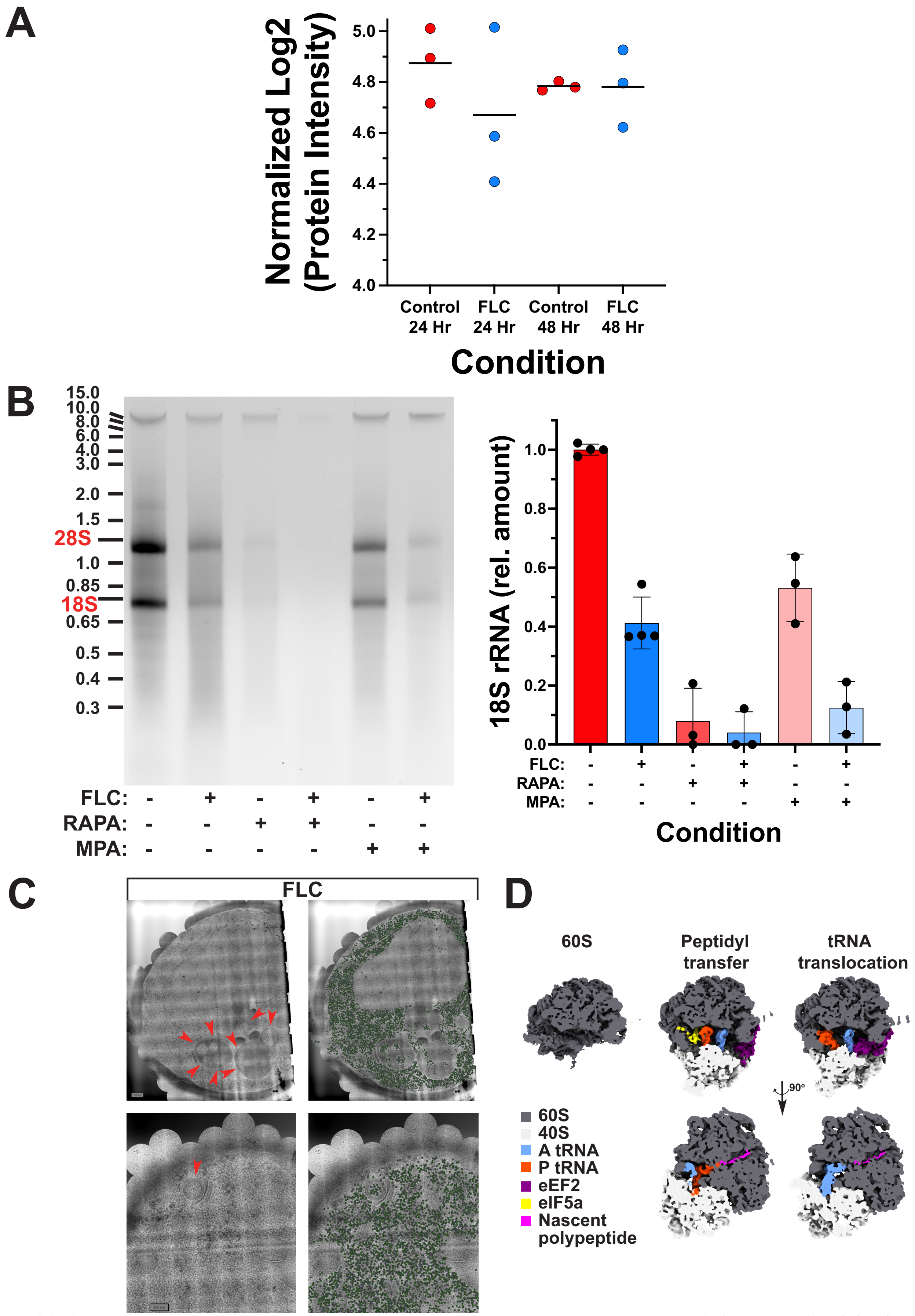


**Figure S6. Ribosomal RNA levels decrease and ribosomal protein levels are constant upon extended growth in fluconazole.** A) Relative abundance of core ribosomal proteins is unaffected by growth in 10X-MIC FLC. Ribosomal proteins levels were determined by Liquid-chromatography tandem mass spectrometry (LC-MS/MS) from independent ($n = 3$) replicates, and normalized to the sample median protein levels. Bars indicate means. B) The rapamycin and mycophenolic acid ribosomal biogenesis inhibitors result in a substantial decrease in rRNA. Total RNA was isolated from the indicated strains and analyzed on an agarose gel (left) and quantitation of 3-4 independent experiments (right) with mean and SD indicated. C) Montage of cryo-EM exposures of two representative FLC-tolerant cells with 60S ribosomal subunit indicated revealing ribosome autophagy. Left images cryo-EM exposures and left images with 60S ribosomal subunit (green) overlays. Red arrow heads indicate regions of autophagy, *i.e.* membrane enclosed structures containing ribosomes. D) Ribosome compositional heterogeneity determined by *in-situ* cryo-EM. Three representative classes of 80S ribosomes corresponding to different stages of translation elongation display compositional heterogeneity. The top view, isosurface rendering, sliced in front of the tRNA binding sites and the bottom view, isosurface rendering, sliced in front of P-site tRNA and ribosome exit tunnel. Color key for displayed components.

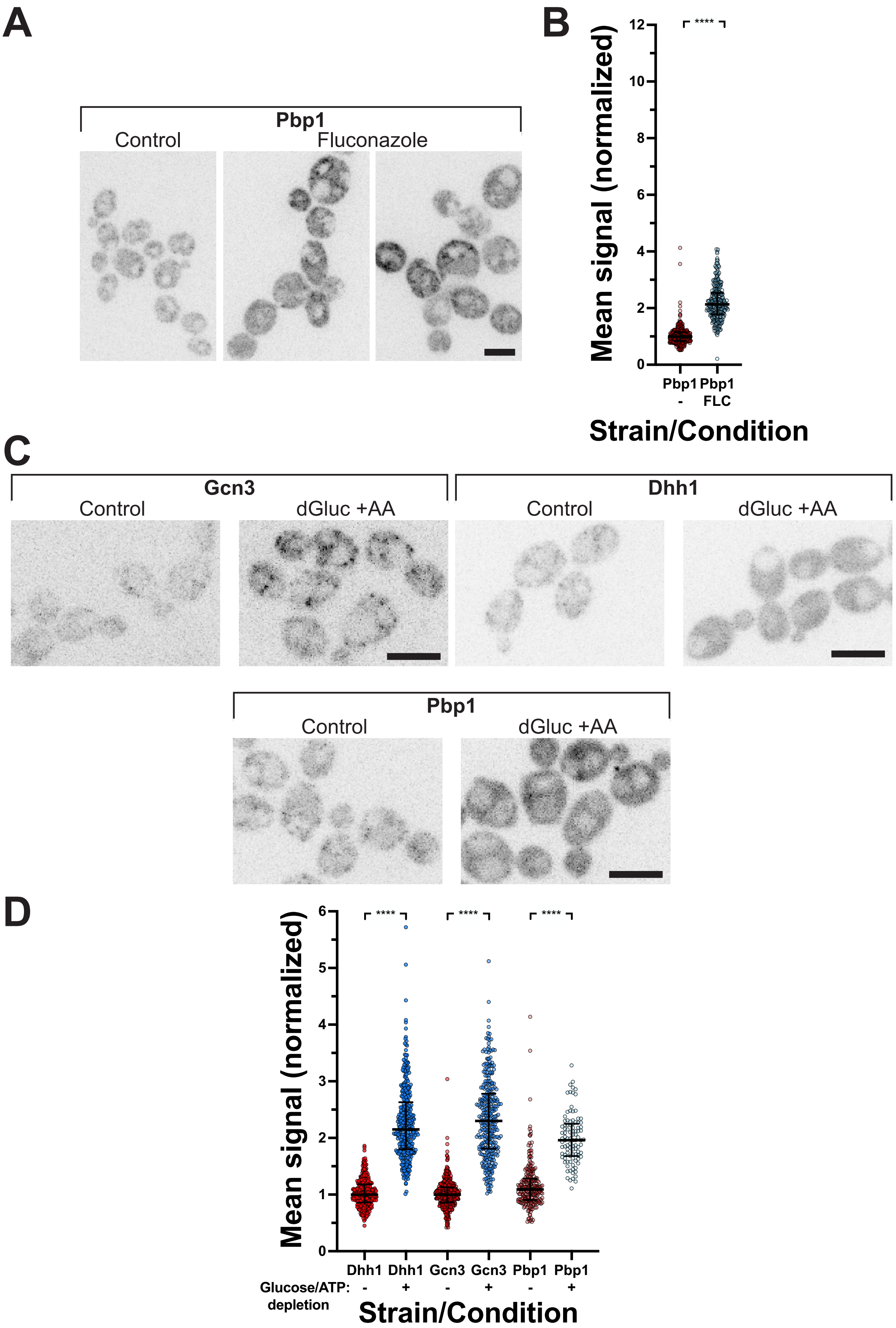


**Figure S7. Growth in fluconazole results two-fold increase stress granule component Pbp1.** A) Pbp1 containing stress granules are evident in the absence and presence of growth in FLC. Images (sum projections of 6 z-sections) of representative cells expressing Ppb1-3xRFP grown in the presence or absence of 10X-MIC FLC for 24 Hr. B) Pbp1 stress granule signal is increased two-fold in cells grown in FLC. Cells treated as in Fig. S7A were used to quantitate signals (*n* = 280-370 cells; from 3 independent experiments), with medians and interquartile ranges indicated with **** < 0.0001). C) Gcn3 condensates are observed upon glucose starvation-induced cytoplasmic acidification. Images (sum projections of 6 z-sections) of representative cells expressing Gcn3-, Dhh1- or Pbp1- 3xRFP grown in the presence or absence of 2-deoxyglucose and antimycin A (GS-ICA conditions; dGluc +AA). Note due to intensity adjustment, intensities between images in Fig. 5F and S7C are not comparable. D) Levels of Dhh1, Gcn3 and Pbp1 increase two-fold upon glucose starvation-induced cytoplasmic acidification. Dhh1, Gcn3 and Pbp1 -3xRFP fusions were used to quantitate signals in cells grown in the presence or absence of GS-ICA conditions (2-deoxyglucose and antimycin A) for 24 Hr (*n* = 100-400 cells; from 3 independent experiments), with medians and interquartile ranges indicated with **** < 0.0001).

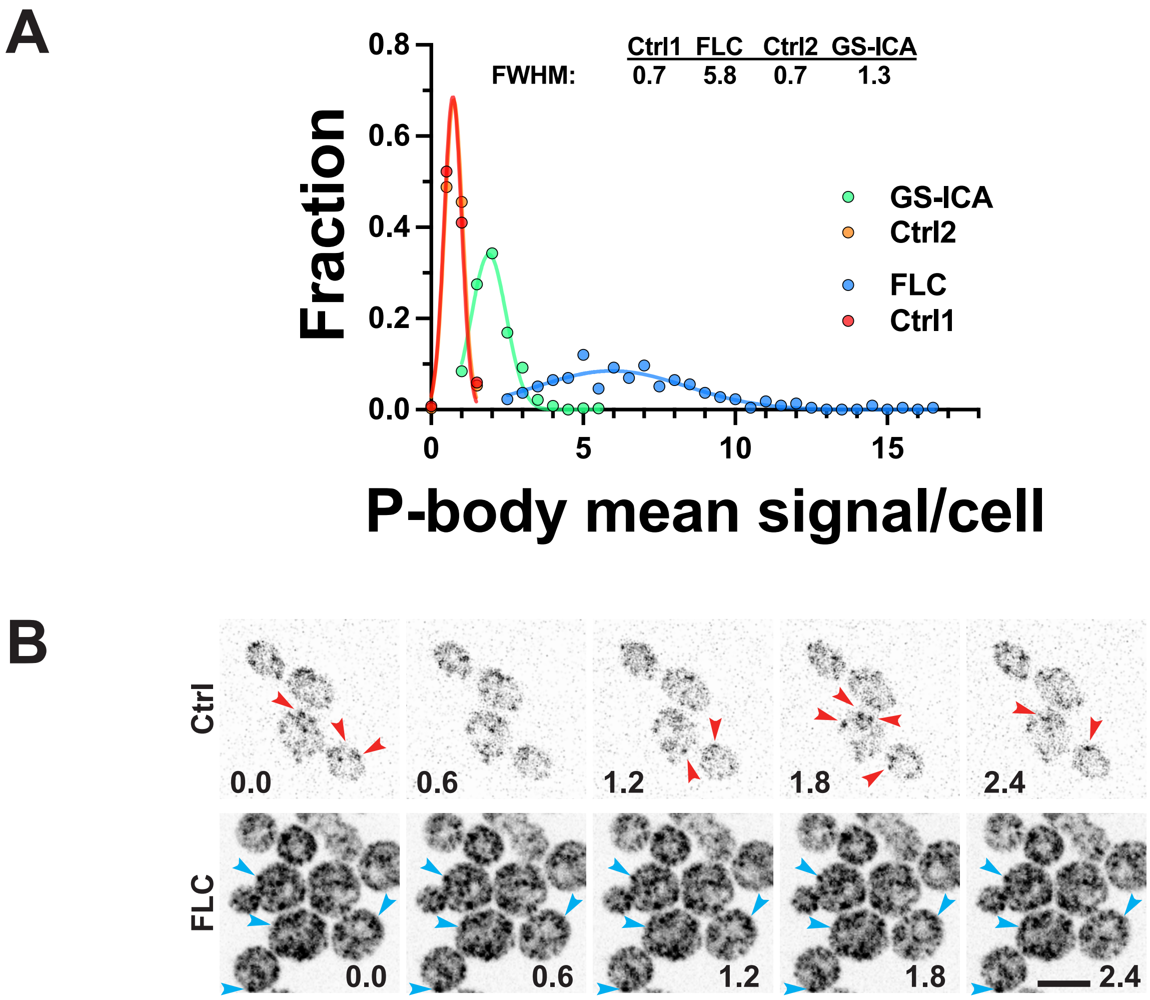


**Figure S8. Processing bodies dynamics are dramatically reduced in fluconazole tolerant cells.** A) Substantial heterogeneity in P-body levels in FLC tolerant cells. A histogram of the mean Dhh1-3xRFP signal per cell from experiments described in Fig. 5C and S7C with bins of normalized (to control conditions) signal of 0.5 is shown. Gaussian curves were fit to the data with $r^2$ values > 0.98 for all conditions except FLC where $r^2$ was 0.85. Full width half maximum (FWHM) for each fitted curve is shown, with an 8.2-fold increase between FLC and control cells and compared to a 1.8-fold between GS-ICA and controls cells. B) Dynamics of P-bodies is altered in cells grown in FLC. Cells expressing Dhh1-3xRFP were imaged and a single Z-section acquired every 0.6 sec. Arrowhead indicate P-bodies (red, control; blue FLC treated).

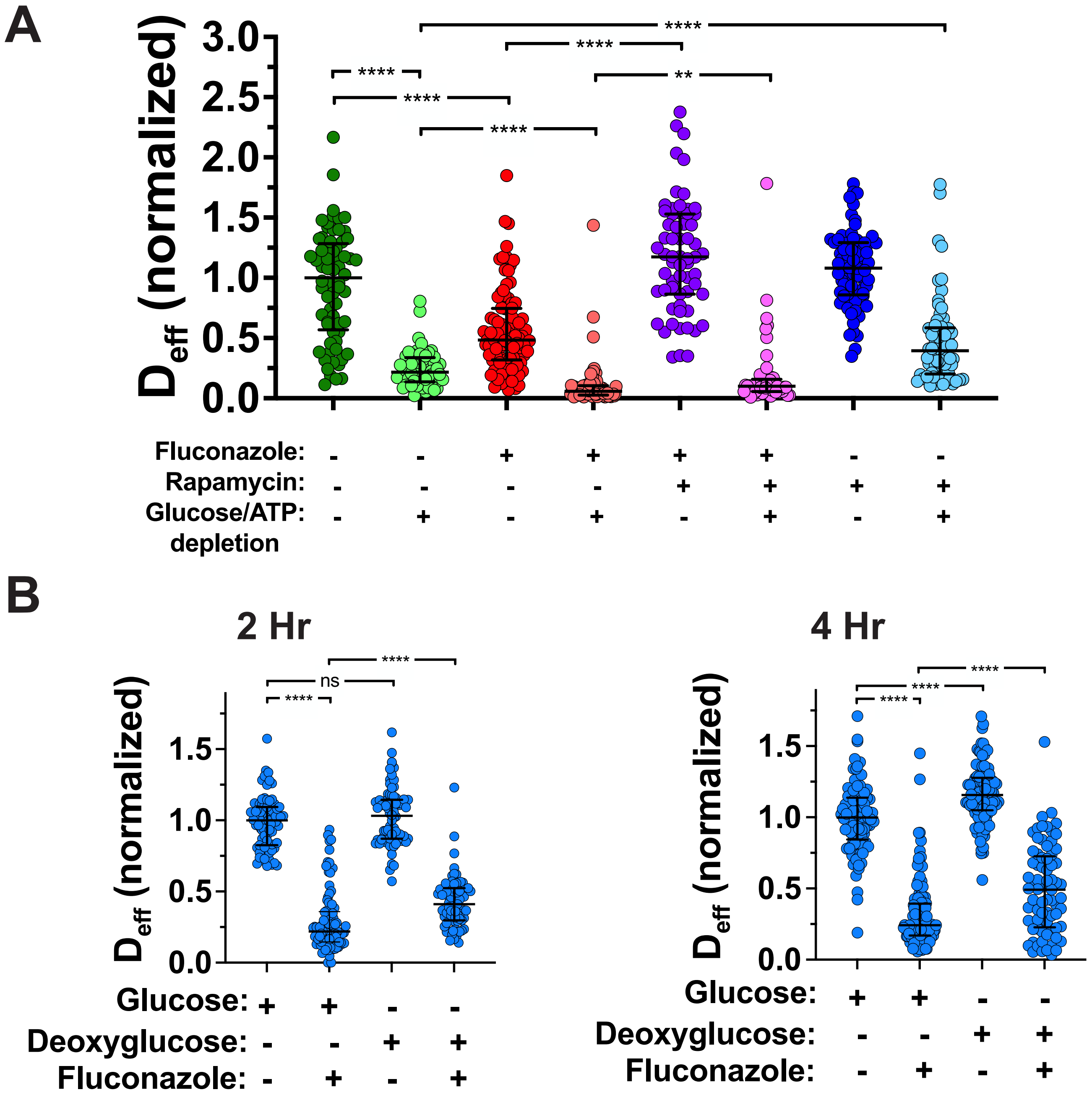


**Figure S9. The fluconazole-dependent decrease in cytoplasmic fluidity is partially restored by glucose-dependent ATP depletion.** A) The FLC-dependent decrease in cytoplasmic fluidity is distinct from glucose starvation-induced acidification the solid-like phase. Cells were grown in the presence of absence of 10X-MIC FLC, rapamycin (RAPA) and/or glucose/ATP depletion for 24 Hr and each symbol is median cell $D_{eff}$ ($n$ = 45-90 cells; 6 - 330 trajectories/cell) normalized to untreated cells. Medians and interquartile ranges indicated with **** < 0.0001 and ** < 0.01. B) Glucose-dependent ATP depletion partially restores the FLC-dependent decrease in cytoplasmic fluidity. Cells were grown in the presence or absence of FLC (10X-MIC) prior to centrifugation and resuspension in either the same media which contained either glucose or 2-deoxyglucose and incubated further for either 2 Hr (left) or 4 Hr (right). Each symbol is median cell $D_{eff}$ ($n$ = 73-124 cells; 25 - 300 trajectories/cell) normalized to untreated cells. Medians and interquartile ranges indicated with **** < 0.0001 and ns not significant.

**Supplementary information**

**Table 1a. Strains used in this study.**

| **Strain number** | **Relevant Genotype** | **Source** |
|---|---|---|
| BWP17 | *ura3Δ::λimm434/ura3Δ::λimm434 his1Δ::hisG/his1Δ::his arg4::hisG/arg4Δ::hisG* | 1 |
| SC5314 | Wild-type reference strain | 2 |
| PY173 | Same as BWP17 with *ENO1/eno1::ENO1-tetR ScHAP4AD-3xHA-ADE2* | 3 |
| PY4865 | Same as BWP17 *RP10::ARG4-ADH1-GFP-CtRac1 adh1Δ::ADH1p-mTrq2-HIS1* | This study |
| PY6414 | Same as BWP17 with *adh1Δ::ADH1p-Pfv-GFPγ*$^{G206K}$*-CdHIS1* | 4 |
| PY6599 | Same as BWP17 with *tef1Δ::TEF1p-Pfv-GFPγ*$^{G206K}$*-CdHIS1* | 4 |
| PY6743 | Same as PY173 with *erg11::URA3pTet*$_{off}$*ERG11/erg11Δ::HIS1* | 5 |
| PY6786 | Same as 6743 with *adh1Δ::ADH1p-Pfv-GFPγ*$^{G206K}$*-ARG4* | This study |
| PY7032 | Same as SC5314 *arg4/arg4 adh1Δ::ADH1p-Pfv-GFPγ*$^{G206K}$*-ARG4* | This study |
| PY7033 | Same as SC5314 *arg4Δ/arg4Δ erg3Δ/erg3Δ adh1Δ::ADH1p-Pfv-GFPγ*$^{G206K}$*-ARG4* | This study |
| PY7034 | Same as SC5314 *arg4Δ/arg4Δ erg3Δ/erg3Δ erg11Δ/erg11Δ adh1Δ::ADH1p-Pfv-GFPγ*$^{G206K}$*-ARG4* | This study |
| PY7072 | Same as 6414 with *GCN3::GCN3-3xmSc-ARG4* | This study |
| PY7301 | Same as PY7173 with *cgr1::URA3pTet*$_{off}$*CGR1/cgr1Δ::CdHIS1* | 4 |
| PY7305 | Same as 6414 with *DHH1::DHH1-3xmSc-ARG4* | This study |
| PY7306 | Same as 6414 with *PBP1::PBP1-3xmSc-ARG4* | This study |
| PY7322 | Same as PY7301with *adh1Δ::ADH1p-Pfv-GFPγ*$^{G206K}$*-ARG4* | 4 |
| PY7406 | with BWP17 with *adh1Δ::ADH1p-mSc-ARG4* | This study |
| PY7546 | Same as BWP17 with *adh1Δ::ADH1p-GFPγ-LoxP-ARG4-LoxP* | 4 |

**Supplementary information**

**Table 1b. Oligonucleotides used in this study.**

| Primer | Sequence (5’ → 3’) |
|---|---|
| CaADH1p-PfVp | CCAGAATTATTTTTTTTTCATCAGTTTAACAACAACAAACGTTATTGTCATACAACAACAACAACAAATACAAAAACAATTATGttatcaattaatccaac |
| CaADH1KixFP_S2 | CTGGGTAATCCTTGTAGACTAATTGACCACCATTGGTATCAAAGACAACGGCTTTTTGAGTTTTTGGGATTTGTTCAGACATtctgatatcatcgatgaattcgag |
| CaADH1KixFP_S1 | CCAGAATTATTTTTTTTTCATCAGTTTAACAACAACAAACGTTATTGTCATACAACAACAACAACAAATACAAAAACAATTatgggtgctggcgcaggtgct |
| Arg4 sgRNA | CGTAAACTATTTTTAATTTGGGTACTAAAGTATACACACAGTTTTAGAGCTAGAAATAGC |
| Erg3 sgRNA | CGTAAACTATTTTTAATTTGgtggttataaagctatcttgGTTTTAGAGCTAGAAATAGC |
| Erg11 sgRNA | CGTAAACTATTTTTAATTTGATTAAACTGAGAAGAGAACGGTTTTAGAGCTAGAAATAGC |
| Donor Arg4-FW | CTTCTGTTCTTTACATAACAGCTATCAAGAATAATCAATTAATAggTTTATAAATAGTCATATAATAATCACAGTATTGTTTAACTTACA |
| Donor Arg4-RV | TGTAAGTTAAACAATACTGTGATTATTATATGACTATTTATAAAccTATTAATTGATTATTCTTGATAGCTGTTATGTAAAGAACAGAAG |
| Donor Erg3-FW | AGTTCAATCTTTTTTTCTTTCTTTCGGATTCGGTTTAGCggTTGGTACATCTTTGTTTTGGACCATTGACTATTTTTCCA |
| Donor Erg3-RV | TGGAAAAATAGTCAATGGTCCAAAACAAAGATGTACCAAccGCTAAACCGAATCCGAAAGAAAGAAAAAAAGATTGAACT |
| Erg11 NAT-FW | TACTTGTCTTCTTTTTATTATATATATAAGTTTCTTTTCAAGAAGATCATAACTCAATtgttatgggtatggatgaattg |
| Erg11 NAT-RV | ACTAAGTAACAAAATGAAAACAATCTGAACACTGAATCGAAAGAAAGTTGCCGTTTTAccgctctagaactagtggatct |
| pSNR52-FW | catctaatcaactcccagat |
| gRNA conserved-RV | taaaaaaaCTCGAGAAAAAAAGCAC |

**Supplementary information**

| | |
|---|---|
| Arg4 up FW | CAAGAGTAGTCTCAAATAAACC |
| Arg4 down-RV | CGTTTGGAAGCTGTATATCG |
| Arg4-internal--FW | TCTTGAACGGGCACATAAAGAAATCG |
| Erg3 up FW | CCATTTCTTTCCCTATTGTGC |
| Erg3 down-RV | CAATATCATCATCACGACCGG |
| Erg3-internal--RV | CTCTGAAAATGTTTGATCTGGC |
| Erg11 up FW | TTGTATTCAATATCGTACCC |
| Erg11 down-RV | TATTAGCAGATGATGCTGGA |
| Erg11-internal--RV | CAGAACCAAACCAAGGAATCC |
| CaGCN3pXFP_S1 | CTCATGAATACATTACTGCTTTGATAACAGATTTGGGAGTATTGA CTCCATCTGCAGTCAGTGAAGAGTTGATAAAAATATGGTACGAT ggtgctggcgcaggtgcttc |
| CaGCN3m_S2 | cacaaactattttgatttgaaaagtttagaaaatgtattagatgaaattatataacaagtccactatatatatatac catcagtattcTCTGATATCATCGATGAATTCGAG |
| CaDHH1pXFP_S1 | GTATCCTCCACAATTTGCTGGCTACCCAGGTCAACCTCCACAACT CCCACAAGGTCAACAACAACATGCTCAAGCACAAAATCCTGCTC AACAATATggtgctggcgcaggtgcttc |
| CaDHH1m_S2 | gggtgcataatcaatagaaaagtaaatgtaaaaataataaaaaaaaaagtaggaaaattatcataaatcatta agtaaagaattgttctacccggaTCTGATATCATCGATGAATTCGAG |
| CaPBP1pXFP_S1 | CGTTTTATCCGCCGCAAGGGTTTGTGCCACCACCCCAGTTTGGTA ACCCGATGATGATGGGCAACCCGTATGGTAGAAGGTACACGAA AAAACATggtgctggcgcaggtgcttc |
| CaPBP1m_S2 | CTGATCTTAGATAACGTTGAATCGCTCATGACGTGGAGAGAGTA AACGAGATTCTTTTTTTGTTGTTCATTTTTTTTTTTTTTCTCTTCC TATATATCTGATATCATCGATGAATTCGAG |
| CaGCN3p827 | GGTTTGTTGATTTCACACC |
| CaDHH1p1329 | GGGTTCATCGAGTCAGCAA |
| CaPBP1p1477 | CCTACGTTTTATCCGCCG |

**Supplementary information**

| | |
|---|---|
| CamScx3Link2m | GCTACCGCTGCCGCTACCCTT |
| ACT1pTM | ATGTTCCCAGGTATTGCTGA |
| ACT1pTM | ACATTTGTGGTGAACAATGG |
| ERG11pTM | TGACCGTTCATTTGCTCAACTATATT |
| ERG11mTM | CACGTTCTCTTCTCAGTTTAATTTCTTTC |

**Supplementary information**
